# Orai1 is required for Ca^2+^-dependent plasma membrane repair and mechanoadaptation

**DOI:** 10.64898/2026.05.13.724989

**Authors:** Haitao Luan, Azize Cerci, Timothy A. Bielecki, Bhopal C. Mohapatra, Santosh Shrestha, Yiyang Wu, Matthew D. Storck, Lynette M. Smith, Kenneth A. Stauderman, Donald W. Coulter, Anupam Kotwal, Jixin Dong, Jung Yul Lim, Vimla Band, Subramanian P. Ramanathan, Hamid Band

## Abstract

Ca^2+^-dependent repair of plasma membrane breaches is essential for animal cell viability. An initial passive influx of extracellular Ca^2+^ triggers the formation of a protein plug that rapidly seals breaches. However, the mechanism of extracellular Ca^2+^ requirement for subsequent repair remains undefined. EHD2 protein stabilizes the plasma membrane caveolae, which sustain membrane repair, and maintains high surface levels of the caveolae-resident Ca^2+^ channel Orai1. We establish the requirement of both Orai1 and EHD2 for repair of plasma membrane lesions induced by mechanical injury or by a model bacterial pore-forming toxin. We demonstrate rapid EHD2 recruitment and Orai1-mediated Ca^2+^ entry at plasma membrane sites of localized mechanical stimulus, the latter requiring EHD2 and CAV1. EHD2 and Orai1 are necessary for mechanosensitive YAP/TAZ-TEAD activation and positive feedback for CAV1 expression that promotes membrane repair. Our studies establish EHD2 and Orai1 as novel components of mammalian plasma membrane repair and mechanoadaptation.

## Introduction

The ability to sense and respond to changes in mechanical properties of the extracellular microenvironment is a fundamental feature of cellular life during development, homeostasis and diseases (Hannezo and Heisenberg 2019, Narain, Muncie-Vasic et al. 2025). Mechanosensing and mechanotransduction are particularly critical for cell types physiologically exposed to dynamic changes in external forces, such as myofibers in skeletal and cardiac muscle exposed to contraction-relaxation cycles, endothelial cells exposed to stretch and shear forces from changing pressure and flow, and lung alveolar cells exposed to cyclical compression during respiration (Martino, Perestrelo et al. 2018). Metastatic tumor cells must similarly navigate through stiff microenvironments during extravasation from the primary tumor, intravasation at metastatic sites and while exposed to flow-associated shear forces in circulation (Narain, Muncie-Vasic et al. 2025). Adaptive mechanisms that allow such cell types to withstand harsh mechanical environments and swiftly repair damaged plasma membrane are therefore of significant interest in understanding physiological systems and pathological states.

Cell types physiologically exposed to high mechanical stress to the plasma membrane, including skeletal and cardiac myocytes, endothelial cells and fibroblasts, show an abundance of caveolae, 50-80 nm diameter cholesterol- and glycosphingolipid-rich plasma membrane invaginations scaffolded on the cytoplasmic side by Caveolin and Cavin proteins (Parton 2018, Sotodosos-Alonso, Pulgarin-Alfaro et al. 2023). Studies using applied external forces and hypo-osmotic conditions have shown a key role for caveolae in protecting the plasma membrane from mechanical force-induced breaches and their repair (Sinha, Koster et al. 2011). Caveolae-mediated endocytosis also plays a key role in repairing plasma membrane pores formed by bacterial pore-forming toxins, by facilitating their lysosomal degradation (Corrotte, Fernandes et al. 2012, Corrotte, Almeida et al. 2013). More recent studies have validated the physiological mechanoprotective role of caveolae in vivo. CAV1 knockout studies show that intact caveolae are required for the integrity of microvascular endothelium, acting against mechanical rupture induced by increased cardiac output (Cheng, Mendoza-Topaz et al. 2015). Loss of caveolae upon Caveolin or Cavin knockdown was also found to cause the collapse of vacuolated cells in Zebrafish notochord under mechanical strain of locomotion (Garcia, Bagwell et al. 2017). Besides their mechanoprotective roles, caveolae function as hubs for cell signaling, ion and nutrient transport, and receptor endocytic traffic (Shvets, Ludwig et al. 2014).

Caveolae respond to plasma membrane tension by flattening to relieve strain (Sinha, Koster et al. 2011), and the related CAV1-containing dolines respond by activating the YAP/TAZ-TEAD pathway (Moreno-Vicente, Pavon et al. 2018, Lolo, Walani et al. 2023). The latter mediate positive feedback through TEAD-dependent induction of CAV1 and CAVIN1 gene expression to sustain caveolae (Dupont, Morsut et al. 2011, Rausch, Bostrom et al. 2019, Lolo, Walani et al. 2023). The contribution of caveolae in the repair of plasma membrane injuries has primarily focused on their role as providers of membrane needed to plug the plasma membrane breaches, either through localized rearrangements or through endocytic/exocytic processes (Sinha, Koster et al. 2011, Corrotte, Fernandes et al. 2012, Corrotte, Almeida et al. 2013, Cheng, Mendoza-Topaz et al. 2015, Garcia, Bagwell et al. 2017, Stefl, Takamiya et al. 2024). In contrast, any roles of caveolae in regulating signaling mechanisms critical for membrane repair are not well-defined.

While Caveolins and Cavins are the structurally required elements of caveolae, accessory proteins localized to caveolae are known to regulate their dynamics at the plasma membrane (Parton 2018, Sotodosos-Alonso, Pulgarin-Alfaro et al. 2023) and thus are candidates to modulate caveolae function in mechanosensing and mechanoprotection. The EPS15 Homology Domain containing protein 2 (EHD2) localizes to and stabilizes the plasma membrane caveolae (Moren, Shah et al. 2012, Stoeber, Stoeck et al. 2012, Hoernke, Mohan et al. 2017, Yeow, Howard et al. 2017), and has been shown to rapidly accumulate at sites of laser-induced plasma membrane lesions in skeletal muscle models, localizing to the shoulder region of the membrane repair cap together with dysferlin (DYSF) (Marg, Schoewel et al. 2012, Demonbreun, Quattrocelli et al. 2016). DYSF and its related family members are known to be involved in plasma membrane repair (Demonbreun and McNally 2016). EHD2 and DYSF interact physically and were required for myotube fusion (Posey, Pytel et al. 2011). Notably, hypotonic stress-induced increase in plasma membrane tension was found to induce rapid translocation of EHD2 into the nucleus and transcriptomic analyses identified EHD2-dependent changes in gene expression (Torrino, Shen et al. 2018), supporting a mechanosensitive role of EHD2. Further, NIH-3T3 cells with combined EHD1, 2 and 4 KO, which reduced the cell surface caveolae reservoir, were found to be vulnerable to PM rupture upon prolonged cyclical stretch (Yeow, Howard et al. 2017). Together, these findings suggest that EHD2 may play a functional role in plasma membrane repair. Whether EHD2 is indeed involved in plasma membrane repair and the mechanisms of such a role are currently unknown.

Our previous studies of the functional role of EHD2 in breast cancer revealed that its overexpression, found in ∼40% of all patients and a majority of HER2+ and triple-negative (TNBC) subtypes, is associated with shorter patient survival and propensity for metastasis (Luan, Bielecki et al. 2023). Knockdown and knockout analyses in TNBC cell models established a pro-tumorigenic and pro-metastatic role of EHD2 (Luan, Bielecki et al. 2023). Mechanistically, we showed that EHD2 was critical to sustain high plasma membrane levels of Orai1 (Luan, Bielecki et al. 2023), a Ca^2+^ channel required for store-operated calcium entry (SOCE). SOCE is a conserved molecular process in which the endoplasmic reticulum (ER) Ca^2+^ depletion induces a conformational change in ER Ca^2+^ sensor STIM1 to promote its translocation to the ER-plasma membrane (ER-PM) contact sites where it binds to and activates Orai1 (Ong, Subedi et al. 2019, Lewis 2020). Orai1-mediated Ca^2+^ entry promotes Ca^2+^-dependent signaling and helps refill the depleted ER stores to protect against unfolded protein response (Elaib, Saller et al. 2016, van Vliet, Giordano et al. 2017). Orai1 is known to reside in and functionally require the cholesterol-rich and CAV1-containing plasma membrane microdomains (Sathish, Abcejo et al. 2012, Chantôme, Potier-Cartereau et al. 2013, Jardin and Rosado 2016, Bohorquez-Hernandez, Gratton et al. 2017). However, whether Orai1-mediated Ca^2+^ entry has any role in caveolae-dependent mechanosensing or mechanoprotection is unknown.

A potential role of the EHD2-Orai1 axis in caveolae-dependent mechanoprotection is a question of broad interest as plasma membrane repair across species is well-established to require extracellular Ca^2+^ entry and to be carried out by Ca^2+^-dependent proteins (Cheng, Zhang et al. 2015, Cooper and McNeil 2015, Demonbreun and McNally 2016, Andrews and Corrotte 2018, Horn and Jaiswal 2018). Based on studies in model organisms, it is widely accepted that the repair process is initiated by passive flow of Ca^2+^ through the breached plasma membrane down the steep concentration gradient from the extracellular space (millimolar Ca^2+^) to the cytoplasm (sub-micromolar Ca^2+^) (Cheng, Zhang et al. 2015, Cooper and McNeil 2015, Demonbreun and McNally 2016, Andrews and Corrotte 2018, Horn and Jaiswal 2018). It is also well-established that the initial sealing of plasma membrane injuries occurs within seconds, but that subsequent repair, which lasts for an extended duration, continues to be Ca^2+^-dependent even though the passive flow of extracellular Ca^2+^ has ceased (Cheng, Zhang et al. 2015, Cooper and McNeil 2015, Demonbreun and McNally 2016, Andrews and Corrotte 2018, Horn and Jaiswal 2018). These late repair steps include the essential roles of Ca^2+^-dependent proteins and Ca^2+^-dependent movement of exocytic and endocytic membrane vesicles that help restore the plasma membrane. The sources of Ca^2+^ required to complete the membrane repair after initial sealing of the plasma membrane lesions remain unclear. The lysosomal Ca^2+^ channel MCOLN1 was found to be important for plasma membrane repair, yet effective repair still required extracellular Ca^2+^, possibly to replenish lysosomal stores (Cheng, Zhang et al. 2014). It is unknown if plasma membrane-localized Ca^2+^ channels contribute to plasma membrane repair.

Recent studies have shown the importance of proteins identified as critical for plasma membrane repair in other models, such as myoferlin (Leung, Yu et al. 2013), annexins (Bouvet, Ros et al. 2020, Gounou, Bouvet et al. 2023) and annexin-associated S100 family members (Jaiswal, Lauritzen et al. 2014), in plasma membrane repair in tumor cells. Notably, the acquisition of a more robust plasma membrane repair capacity was identified as a response to the higher propensity of invasive breast cancer cells to undergo increased plasma membrane damage (Jaiswal, Lauritzen et al. 2014), supporting the idea that plasma membrane repair in cancer cells represents a functionally important mechanoprotective adaptation. As we linked the EHD2-Orai1 axis to the stability of CAV1-containing plasma membrane domains and invasive/metastatic behavior of TNBC cells (Luan, Bielecki et al. 2023), we utilized these cell models to examine the role of the EHD2-Orai1 axis in caveolae-dependent mechanoprotection.

Our findings establish that Orai1 and its ability to import the extracellular Ca^2+^ into cytoplasm are required for the repair of plasma membrane injuries induced mechanically or by a model bacterial pore-forming toxin, streptolysin O (SLO). We demonstrate that mechanical force applied to the plasma membrane elicits rapid, highly localized, EHD2 recruitment and Orai1-mediated Ca^2+^ influx, identifying the mechanosensitive nature of the EHD2-Orai1 axis. Finally, we show that stiff extracellular matrix activation of YAP/TAZ-TEAD signaling requires the EHD2-Orai1 axis-dependent Ca^2+^ import, and in turn helps sustain Orai1-mediated mechanosensing and mechanoprotection. Thus, our studies establish a new paradigm for plasma-membrane-channel-mediated import of extracellular Ca^2+^ as an essential component of mammalian plasma membrane repair and identify a novel mechanosensitive role for EHD2 and Orai1 in signaling to the YAP/TAZ pathway.

## Results

### EHD2 is required for efficient repair of plasma membrane injuries induced by mechanical rupture or streptolysin O

In view of the established role of EHD2 to stabilize the plasma membrane pool of caveolae (Moren, Shah et al. 2012, Stoeber, Stoeck et al. 2012, Hoernke, Mohan et al. 2017, Yeow, Howard et al. 2017), recruitment of EHD2 to the shoulder region of plasma membrane repair cap in skeletal muscle injury models (Marg, Schoewel et al. 2012, Demonbreun, Quattrocelli et al. 2016), mechanical perturbation-induced nuclear shuttling of EHD2 and its involvement in gene expression (Torrino, Shen et al. 2018), we posited that EHD2 may be required for plasma membrane repair. We adapted the scratch wounding protocol commonly used to assess tumor cell migration to examine mechanically induced plasma membrane injury repair since a large proportion of cells near the scratch wound border showed the uptake of membrane impermeant fluorescent dyes, indicative of cells with plasma membrane damage (**Fig. 1A, Fig. S1A & S1B**). Incubation of wildtype (WT) MDA-MB231 or Hs578T TNBC cell lines for various time points in Ca^2+^-containing medium demonstrated that most cells that incorporated the membrane-impermeant fluorescent dye FITC-dextran (i.e., cells with plasma membrane injury) became impermeant to the subsequently added propidium iodide (PI) within 1-5 minutes with slower recovery after that (**Fig.1B**), indicating successful plasma membrane repair. In contrast, the injured cells incubated in medium without Ca^2+^ showed significantly impaired repair (70% vs. 40% cells with repair at 40 min in +Ca^2+^ vs. -Ca^2+^ media; p<0.05) (**Fig. 1B**). Thus, as expected, the TNBC cell models we use exhibit robust Ca^2+^-dependent plasma membrane repair. Compared to WT TNBC cells, their EHD2-KO versions exhibited a marked and significant reduction in the levels of plasma membrane repair (25% vs. 70% repair at 40 min, p<0.01), close to that observed in WT cells in the absence of Ca^2+^; absence of Ca^2+^ further reduced the plasma membrane repair of EHD2-KO cells, but the difference was smaller (20% vs. 15% in +Ca^2+^ vs. -Ca^2+^, p>0.05) (**Fig. 1B**). To further explore the role of EHD2 in plasma membrane repair, we used streptolysin O (SLO) to induce plasma membrane pores. SLO is a prototype bacterial toxin that forms smaller and more uniform plasma membrane pores that are also repaired in a Ca^2+^-dependent process (Cheng, Zhang et al. 2015, Cooper and McNeil 2015, Demonbreun and McNally 2016, Andrews and Corrotte 2018, Horn and Jaiswal 2018). Repair was assessed by analyzing the proportion of cells permeable to PI using FACS analysis. In contrast to WT TNBC cells, EHD2-KO cells exhibited a significantly higher percentage of PI-high cells (40% vs. 16% PI+ cells in MDA-MB-231 and 60% vs. 35% PI+ cells in Hs578T; p<0.001), indicating less efficient repair (**Fig. 1C &1D**). Notably, EHD2-KO MDA-MB231 cells reconstituted with mouse EHD2 (Luan, Bielecki et al. 2023) showed repair comparable to that in WT cells (20% vs. 16% PI+ cells; p>0.05) (**Fig. 1C &1D**). Together, these results led us to conclude that EHD2 is required for the repair of plasma membrane injuries induced by mechanical force or a prototype pore-forming bacterial toxin.

**Figure 1.**
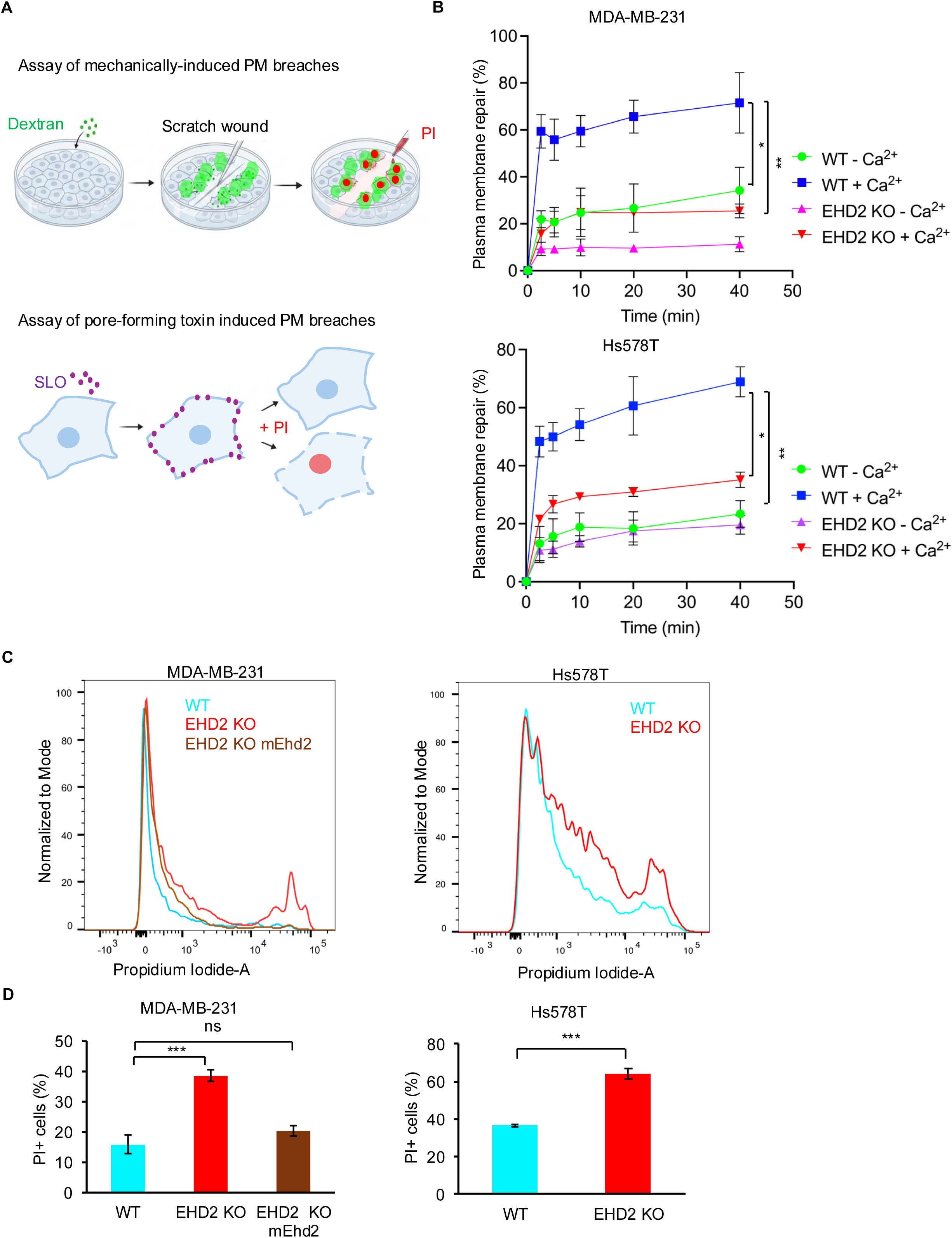
EHD2 is required for plasma membrane repair. **A-B.** Demonstration of EHD2 requirement for mechanically-induced Ca^2+^-dependent plasma membrane repair in TNBC cell lines. The indicated cell lines were subjected to cell scraping to induce mechanical injury to plasma membrane in media containing FITC-dextran (to label cell with damaged plasma membrane) with or without Ca^2+^. At the indicated times after wounding at room temperature, cells were rinsed and incubated with propidium iodide to label cells that had failed to repair. The number of wounded cells with successful repair (only FITC-Dextran-labeled) are shown as a percentage of total cells (green cells plus green and red cells). **A.** Schematic diagrams of plasma membrane repair assays induced by mechanical injury or Streptolysin O toxin**. B.** Quantification of cells with successful plasma membrane repair over time. Data represents mean +/- SEM of three experiments, two-way ANOVA, *,p<0.05, **,p<0.01. **C-D.** Demonstration of EHD2 requirement for the repair of streptolysin O (SLO) induced plasma membrane pores. The indicated cell lines were treated with SLO for 5 min in Ca^2+^ free Tyrode’s buffer on ice, followed by incubation for 10 min in Ca^2+^-containing Tyrode’s buffer. Cells were incubated with propidium iodide-containing medium and analyzed by FACS. **C.** Representative FACS analyses of propidium Iodide (PI) staining after streptolysin O (SLO)-induced membrane damage and repair. Cells to the right of the main peak on left represent those that failed to repair their plasma membrane. **D.** Quantification of PI positive cell population shown in (C). EHD2 KO-mEhd2 represents EHD2-KO MDA-MB-231 cells rescued by stable expression of mouse Ehd2. Quantified data shown are from three independent experiments. Welch’s t-test. *,p<0.05, **,p<0.01, ***, p<0.001.

### Plasma membrane repair is mediated by the activity of Orai1 Ca^2+^ channel

Previously, we showed that loss of EHD2 expression leads to lower plasma membrane levels of Orai1, and the functional impact of the EHD2 loss on cell migration and tumorigenesis was recapitulated by Orai1 inhibition while overexpression of STIM1 partially rescued the SOCE and cell migration defects in EHD2-KO MDA-MB-231 cells (Luan, Bielecki et al. 2023). We therefore tested the possibility that Orai1 may be required for EHD2-dependent plasma membrane repair. First, we generated Orai1-KO derivatives of MDA-MB-231 and Hs578T TNBC cell lines and confirmed the absence of Orai1 protein expression (**Fig. 2A**). Similar to our previous findings upon EHD2-KO in these cell models (Luan, Bielecki et al. 2023), the extent of initial Ca^2+^ release as a measure of ER Ca^2+^ stores (50 % reduction in Orai1-KO in MDA-MB-231, p<0.001; 43 % reduction in Orai1-KO in Hs578T, p<0.005) and their SOCE response to Ca^2+^ store depletion induced by thapsigargin (∼65% reduction in Orai1-KO, p<0.001) (**Fig. 2B& 2C**), as well as their trans-well cell migration towards serum-containing medium (∼60% reduction in Orai1-KO, p<0.001) (**Fig. S2A**), were markedly and significantly reduced. The reduction in ER release reflects a deficit of ER Ca^2+^ store filling because of impaired SOCE (Luan, Bielecki et al. 2023). Notably, Orai1-KO led to a significant impairment in the repair of mechanically-induced plasma membrane injuries (65% in WT vs. 38% in KO MDA-MB-231; 62% in in WT vs. 40% KO Hs578T at 40 min, p<0.01) (**Fig. 2D& 2E, Fig. S2B**) and SLO-induced membrane pores (14% PI^+^ cells in WT vs. 36% in KO in MDA-MB-231 and 19% PI^+^ cells in WT vs. 48% in KO in Hs578T; p<0.001) (**Fig. 2F& 2G**). Complementing the genetic approach, we also assessed the impact of Orai1 inhibition. As the tool inhibitors used in previous studies, such as SKF-96365, lack selectivity (Ramsey, Delling et al. 2006, Ding, Zhang et al. 2012), we utilized a more recently developed Orai1-selective inhibitor CM4620 which functions by inhibiting the activated state of Orai1 (Stauderman 2018, Waldron, Chen et al. 2019) and has progressed through phase 2 clinical trials against acute pancreatitis and COVID-19 pneumonia (Miller, Bruen et al. 2020, Bruen, Miller et al. 2021, Bruen, Al-Saadi et al. 2022). First, we established the Orai1 dependence of the CM4620 effect on functional readouts of Orai1 activity in TNBC cells. Indeed, CM4620 robustly inhibited the SOCE (∼67% reduction in CM4620-treated vs. control; p<0.001) and cell migration (∼55% reduction with CM4620 vs. control; p<0.001) in WT TNBC cells but had little impact on the residual cell migration in Orai1-KO cells (185 cells per field with DMSO vs. 180 cells per field with CM4620, not significant) (**Fig. 2H** & **Fig. S2D**). Importantly, treatment with CM4620 impaired the repair of mechanical (42% in CM4620-treated cells vs. 74% in control cells, p<0.01) (**Fig. 2I & Fig. S2E**) as well as SLO-induced (38% PI+ cells in CM4620 vs. 12% PI+ cells in control, p<0.001) (**Fig. 2J**) plasma membrane damage in TNBC cells. Together, these results support the conclusion that plasma membrane Ca^2+^ channel Orai1 is required for efficient plasma membrane repair.

**Figure 2.**
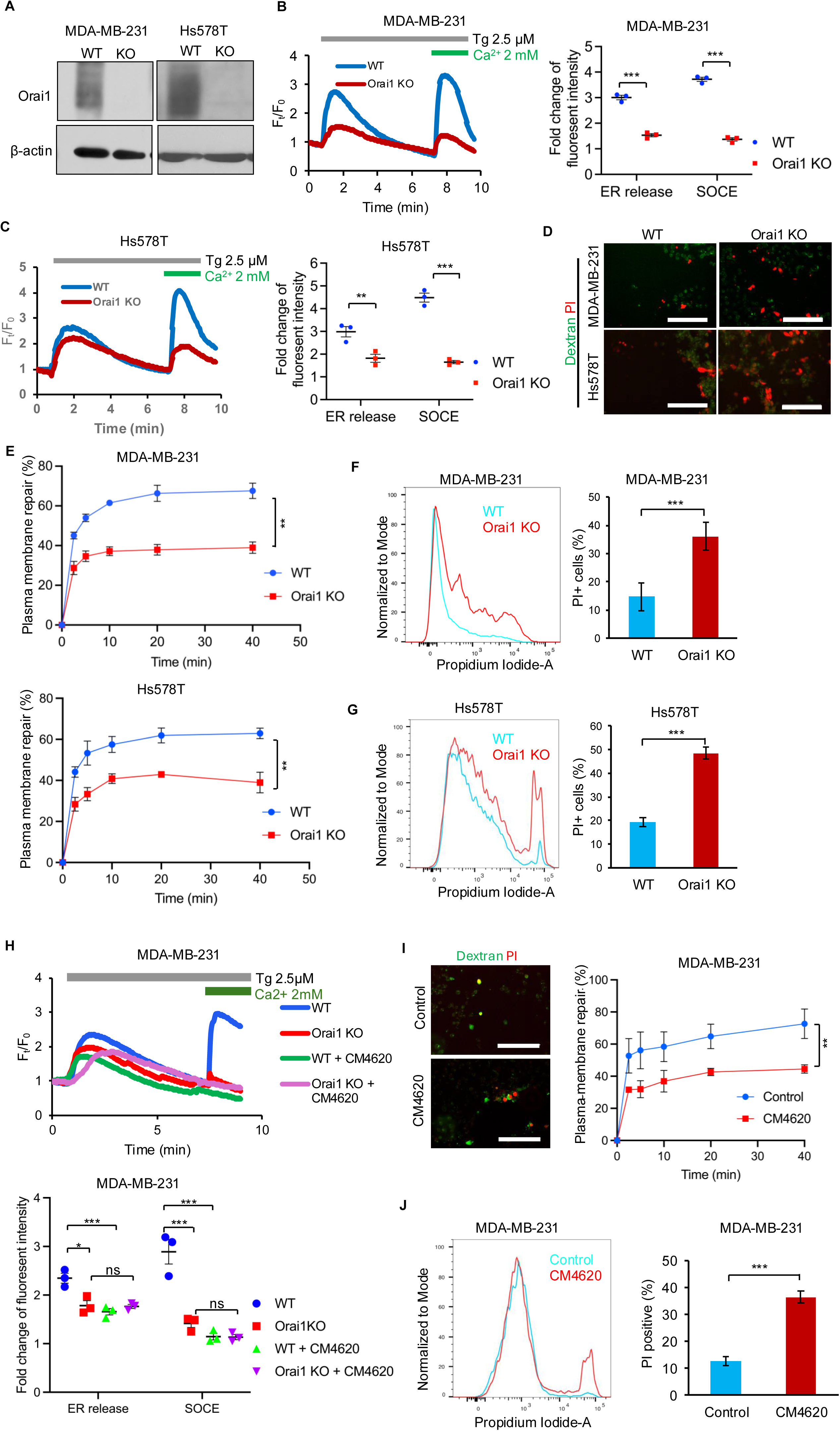
Orai1 is required for plasma membrane repair. **A.** Western blot confirmation of loss of Orai1 expression upon CRISPR/Cas9-mediated KO in MDA-MB-231 and Hs578T cells; shown are pools of three clones maintained individually. **B-C.** Impairment of SOCE upon Orai1 KO in TNBC cells. SOCE was measured using thapsigargin-induced ER Ca^2+^ depletion. Typical profiles are shown on left and quantification of fold-change in peak fluorescence intensity from 3 independent experiments is shown on right. Welch’s t test, ***p<0.001. **D-E.** Orai1-KO impairs the repair of mechanically induced plasma membrane injury. WT vs. Orai1-KO cells were subjected to mechanical injury to plasma membrane and repair assay was performed as in Fig. 1A-B. Representative confocal images are shown in D and quantified data are shown in E. Scale bar, 200 µm. Data represents mean +/- SEM of three experiments, two-way ANOVA, **,p<0.01. **F-G.** Orai1-KO impairs the repair of SLO-induced membrane injury. Plasma membrane damage using SLO and the repair assay on MDA-MB-231 (**F**) and Hs578T (**G**) cell lines were as in Fig. 1C-D. Representative confocal images are shown on left. Quantification of PI-stained cells from 3 independent experiments is shown on right. Welch’s t test, ***p<0.001. **H-J.** Orai1 inhibitors impair plasma membrane repair. Panel **H** shows Thapsigargin (TG)-induced SOCE measurements (initial peak, Ca^2+^ store release in the absence of extracellular Ca^2+^; second peak, SOCE in the presence of extracellular Ca^2+^) performed on the indicated cell lines cultured with or without CM4620 (10 μM, 4 hours pretreatment). Note significant SOCE inhibition by CM4620 in WT cells but not in Orai1-KO cells, supporting Orai1-selelctive effect of CM4620. Data represents mean +/- SEM of three experiments, Welch’s t test, ***p<0.001. The **I** panel shows the impairment of mechanically induced plasma membrane repair by CM4620 in MDA-MB-231. Cells were cultured without or with pretreatment with CM4620 (10 μM) and analyzed for repair as in Fig. 1A-B. Scale bar, 200 µm. Data represents mean +/- SEM of three experiments, two-way ANOVA, **,p<0.01. The **J** panel shows the impairment of SLO-induced membrane damage in MDA-MB-231. Left panel shows representative FACS analysis of membrane repair in cells without or with CM4620 treatment. Right panel shows quantification of cells that failed to repair (PI staining) from 3 independent experiments. Data represents mean +/-SEM of three experiments. Welch’s t test, ***, p<0.001.

### EHD2 and Orai1 are required for mechanosensitive spatiotemporally regulated import of calcium into the cytoplasm

The requirement of EHD2 and Orai1 for plasma membrane repair suggested that these proteins orchestrate a novel pathway of mechanosensitive entry of Ca^2+^ from the extracellular space into the cytoplasm. To test this possibility, we used Atomic Force Microscopy (AFM) to assess the impact of a mechanical stimulus applied to the plasma membrane. MDA-MB-231 cells transfected with fluorescent EHD2 were subjected to nano-indentation with cantilevers and localization of EHD2 over time was monitored by confocal imaging (**Fig. 3A**). We observed rapid (within seconds) accumulation of fluorescent EHD2 precisely at the indentation site (indentation point indicated), which dissipated quickly once the mechanical force was removed (**Fig. 3B & 3C**). The focal accumulation of EHD2 signals was significantly higher compared to the pre-induction signals (∼2 fold higher at the peak, p<0.01) (**Fig. 3D & 3E**). Next, we used MDA-MB-231 cells transfected with a red fluorescent reporter of cytoplasmic Ca^2+^ (R-GECO1.2) (Wu, Liu et al. 2013) to assess if the localized application of force to the plasma membrane induced Ca^2+^ import into cytoplasm. We observed rapid Ca^2+^ entry that started near the site of the plasma membrane indentation and spread to rest of the cell; the Ca^2+^ entry dissipated quickly when the mechanical stimulus was removed (**Fig. 4A**). The mechanosensitive Ca^2+^ entry was abrogated by genetic KO of EHD2 (∼45% reduction; p<0.01), Orai1 (∼40% reduction; p<0.01) or CAV1 (∼55% reduction; p<0.01), and the defect in EHD2-KO cells was partially rescued by ectopic expression of mouse *Ehd2* (∼45% reduction in EHD2-KO vs. 20% reduction in m*Ehd2*-reconstituted cells; p<0.05) (**Fig. 4B**). Further, an Orai1 inhibitor CM5480 (Stauderman 2018, Pallagi, Görög et al. 2022, Szabó, Csákány-Papp et al. 2023), which also exhibited Orai1-dependent activity in TNBC cells (**Fig. S3A**), effectively inhibited the mechanosensitive Ca^2+^ entry (∼85% reduction in CM5480 vs. control, p<0.01) (**Fig. 4C**). To more directly interrogate if the mechanosensitive, Orai1-dependent, Ca^2+^ import observed above indeed reported the Orai1-mediated Ca^2+^ entry, we transfected MDA-MB231 with a genetically encoded fluorescent biosensor, G-GECO1-Orai1 (Dynes, Amcheslavsky et al. 2016). In this biosensor, the green-fluorescent Ca^2+^ indicator fused to the N-terminus of Orai1 itself detects the Orai1-associated Ca^2+^ influx locally in the cytoplasmic nanodomain adjacent to the plasma membrane (Dynes, Amcheslavsky et al. 2016). Transiently transfected G-GECO1-Orai1 showed plasma membrane localization as expected (**Fig. 4D**). The G-GECO1-Orai1 also accurately reported only the SOCE phase of the Ca^2+^ influx in response to thapsigargin treatment of MDA-MB-231 cells, which was completely abolished by the selective Orai1 inhibitor CM5480 (**Fig. S3B**). Indentation of the plasma membrane led to marked, and statistically-significant, increase in Ca^2+^ influx reported by G-GECO1-Orai1 (6.2-fold increase in signal over unstimulated cells; p<0.001) (**Fig. 4D**). Pretreatment of cells with CM5480 led to a highly significant inhibition of G-GECO1-Orai1 fluorescence upon indentation (1-fold-change in CM5480-treated cells vs. 6.2-fold change in control; p<0.001) (**Fig. 4D**). These results conclusively establish that mechanical force applied to the plasma membrane leads to Orai1 activation. Since Orai Ca^2+^ channels are gated by STIM proteins (Ong, Subedi et al. 2019, Lewis 2020), we asked if STIM proteins are required for mechano-sensitive Orai1 activation. In MDA-MB231 cells, both STIM1 and STIM2 were robustly expressed, and siRNA KD of STIM2 led to a substantial upregulation of STIM1 expression (**Fig. S3C**). Concurrent STIM1 and STIM2 siRNA transfection in MDA-MB231 cells expressing R-GECO1.2 led to efficient knockdown of both STIM1 and 2 (**Fig. S3D**) and effectively abolished the plasma membrane indentation induced Ca^2+^ influx (**Fig. 4E**). These results establish that EHD2-, CAV1- and Orai1-dependent mechano-sensitive Ca^2+^ entry is indeed mediated by Orai1 and dependent on STIM proteins. Altogether, these results identify EHD2 and Orai1 as components of a novel axis that mediates mechanosensitive Ca^2+^ import from the extracellular space into the cytoplasm with high spatial and temporal control.

**Figure 3.**
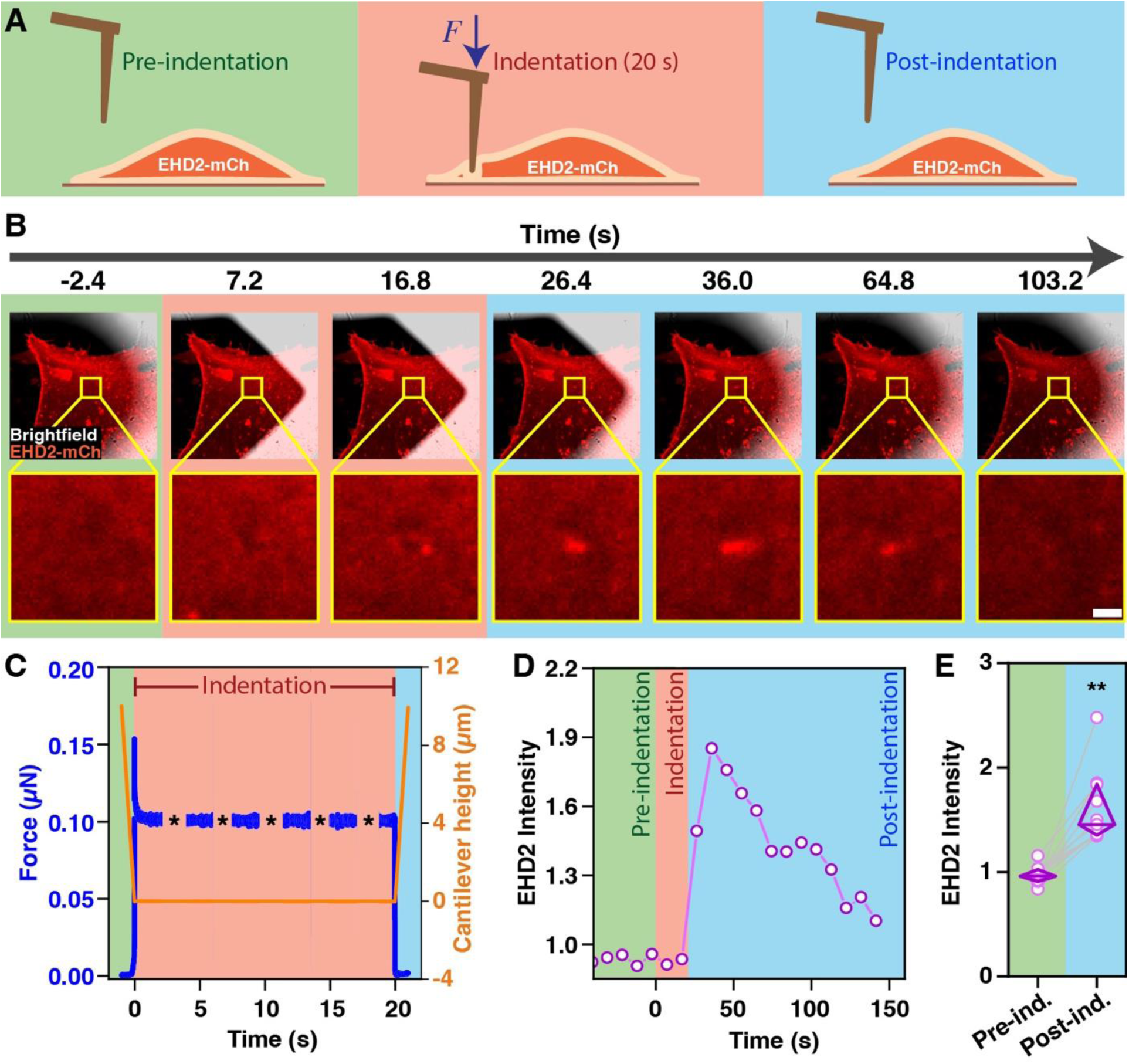
Rapid recruitment of EHD2 to plasma membrane at sites of mechanical force application. **A.** Schematic diagram of force application to plasma membrane using indentation using an atomic force microscope (AFM). MDA-MB231 cells plated on glass-bottom dishes were transiently transfected with EHD2-mCherry (Stoeber, Stoeck et al. 2012) and imaged with a confocal microscope while AFM micro-cantilevers were used to apply force at specific locations on the plasma membrane (indentation). Pre-indentation, green; indentation, red; post-indentation, blue. **B.** Representative time-lapse image of EHD2-mCherry transfected MDA-MB-231 cells at the indicated times before, during and after indentation. The yellow box indicates the area of membrane indentation. Lower panels show higher-magnification images highlighting the area around the indentation. Scale bar, 5 µm. **C.** Measurement of the force applied to the cell (μN; blue) in relation to cantilever indentation (depth in μM, red). **D.** Quantification of EHD2 fluorescence intensity at various time points during plasma membrane indentation. **E.** Fold change of EHD2 fluorescence intensity before and after indentation (1 min). Data points represent cells analyzed through three independent experiments (n= 12). Two-way ANOVA; **p<0.01.

**Figure 4.**
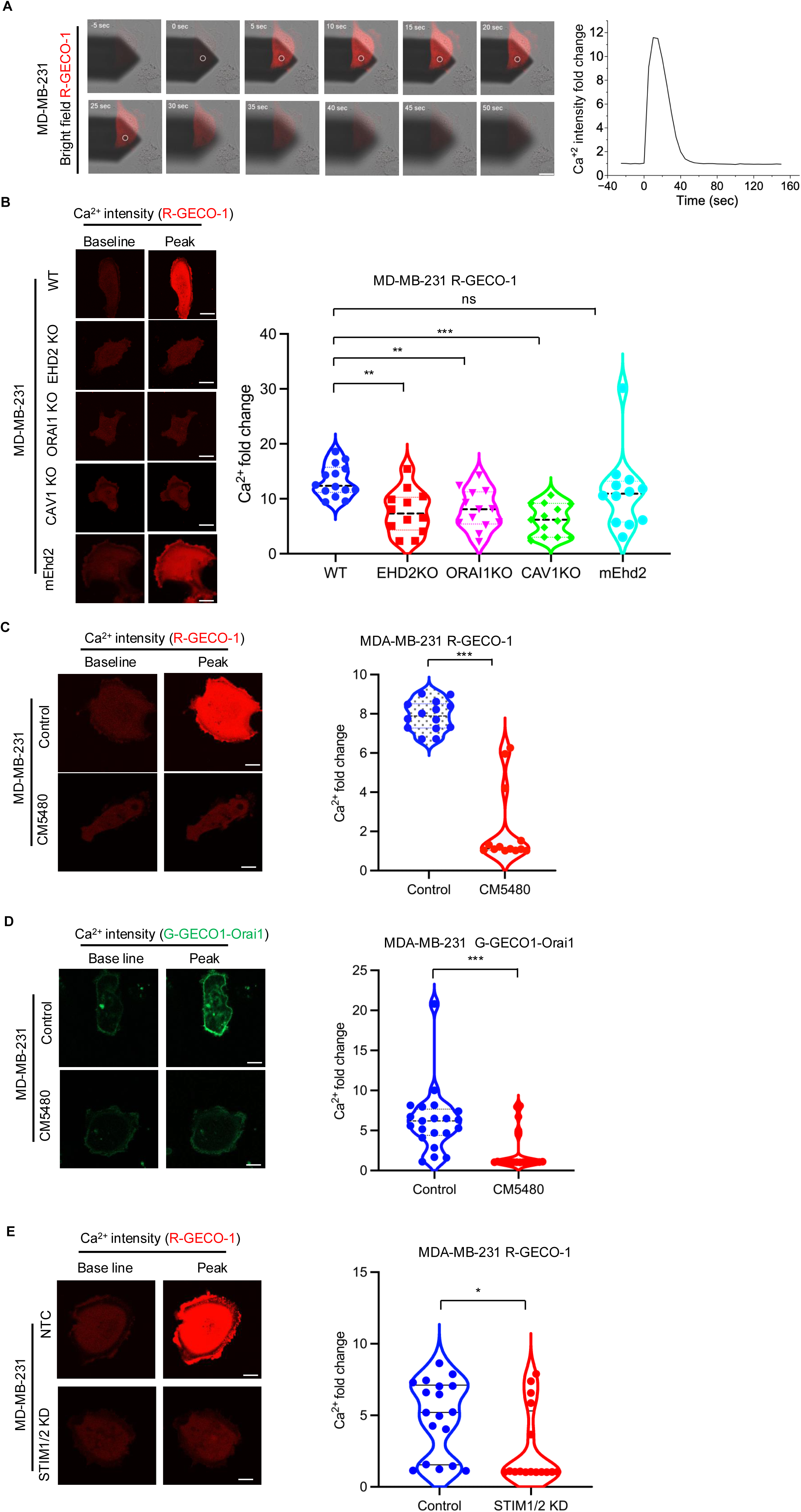
EHD2 and Orai1 are required for mechanosensitive extracellular calcium import into cytoplasm. MDA-MB231 cells plated on glass-bottom dishes were transiently transfected with the Ca^2+^ reporter R-GECO1.2. Cells were imaged with a confocal microscope while micro-cantilevers were employed to indent specific locations on the plasma membrane. **A.** Representative time-lapse images wherein white circles indicate the location of membrane indentation (Left panel). Quantification of Ca^2+^ intensity fold change (Right panel). **B.** Abrogation of mechanical force-induced Ca^2+^ import by EHD2-, Orai1- or CAV1-KO and rescue of EHD2-KO cell response with stable mouse EHD2 (mEHD2) expression. Left, representative images at baseline and peak of AFM indentation. Right, quantification of Ca^2+^ signals. Shown are fold change in Ca^2+^ reporter (R-GECO1.2) signals in MDA-MB-231 cell lines without (WT) or with the indicated genetic perturbations. Fold change of Ca^2+^ peak fluorescence intensity after indentation relative to basal fluorescence intensity prior to indentation are computed from three independent experiments (n=14). One-way ANOVA with Dunnett’s multiple comparisons test, ***p<0.001, **p<0.01; ns, not significant. **C**. Inhibition of mechanical force-induced Ca^2+^ import by Orai1 inhibitor CM5480. Ca^2+^ reporter R-GECO1.2-transfected MDA-MB-231 cells cultured without or with CM5480 (10 μM; 24 h pretreatment) were subjected to AFM cantilever indentation and fluorescence intensity recorded over time. Data points represent cells (n= 14) analyzed through three independent experiments. Welch’s t test, ***p<0.001. **D**. Mechanical force induced Ca^2+^ import recorded by Orai1-linked Ca^2+^ biosensor and its inhibition by CM5480. Orai1-linked Ca^2+^ biosensor (G-GECO1-Orai1)-transfected MDA-MB-231 cells cultured without or with CM5480 (10 μM) were subjected to indentation with AFM cantilever and fluorescence intensity recorded over time. Left, representative images at baseline and peak of indentation. Right, quantification of data. Data points represent cells (n= 21) analyzed through three independent experiments. Welch’s t test, ***p<0.001. **E.** Inhibition of mechanical force-induced Ca^2+^ import upon combined STIM1 and STIM2 siRNA knockdown. MDA-MB-231 cells transfected with the Ca^2+^ reporter (R-GECO1.2) and co-transfected with control or STIM1 and STIM2 siRNAs (knockdown verified in Fig. S3D) were subjected to AFM cantilever indentation and fluorescence intensity recorded over time. Left, representative images at baseline and peak of indentation. Right, quantification of data. Data points represent cells (n=18) analyzed through three independent experiments. Welch’s t test, *p<0.05. Scale bar, 5 µm.

### EHD2-Orai1 axis is required for mechanosensitive activation of the YAP/TAZ-TEAD pathway

Mechanical cell strain, the primary driver of plasma membrane damage, is typically elevated in stiff extracellular microenvironments, including in tumors (Lachowski, Matellan et al. 2022, Lee, Yun et al. 2025). Recent work has established that application of mechanical force to plasma membrane induces CAV1-dependent YAP/TAZ-TEAD pathway activation (Dupont, Morsut et al. 2011, Rausch, Bostrom et al. 2019, Lolo, Walani et al. 2023), and there is emerging support for Ca^2+^ as a potential intermediate to positively or negatively modulate such mechanosensitive YAP/TAZ activation (Wei and Li 2021). Importantly, the downstream targets of mechanosensitive YAP/TAZ-TEAD activation include CAV1 and CAVIN1 in positive feedback that was found to be essential to sustain high levels of plasma membrane caveolae (Dupont, Morsut et al. 2011, Rausch, Bostrom et al. 2019, Lolo, Walani et al. 2023). Our findings that EHD2 and Orai1 are required for mechanosensitive Ca^2+^ entry from the extracellular space into cytoplasm raised the possibility that EHD2-Orai1 axis serves as a mechanosensitive activator of YAP/TAZ signaling. As reported (Moreno-Vicente, Pavon et al. 2018), culture of WT MDA-MB231 cells on stiff matrix (64 kPa) induced the nuclear translocation of YAP compared to cells cultured on soft hydrogel (0.2 kPa) (∼6 fold higher nuclear/cytoplasmic ratio of YAP staining on 64 kPa vs. 0.2 kPa hydrogel, p<0.001) (**Fig. 5A& 5B, Fig. S5**). Analysis of KO cell lines revealed that while YAP nuclear translocation was still significantly higher on stiff compared to soft matrix (nuclear/cytoplasmic YAP ratio ∼1.5 fold higher in EHD2-KO, ∼3 fold higher in Orai1-KO and ∼4 fold higher in Cav1-KO;p<0.001) (**Fig. 5A& 5B, Fig. S5**), the extent of YAP nuclear translocation in EHD2-KO, Orai1-KO and CAV1-KO MDA-MB-231 cells was significantly reduced compared to that in WT cells (∼83% reduction in EHD2-KO vs. WT, ∼ 86% reduction in Orai1-KO vs. WT, ∼ 78% reduction in CAV1-KO vs. WT on 64 kPa matrix; p<0.001) (**Fig. 5A& 5B, Fig. S5**). Notably, mouse *Ehd2* expression in EHD2-KO cells partially restored the YAP nuclear translocation (∼3.5-fold increase in mouse *Ehd2*-rescued vs. 1.5-fold increase in EHD2-KO cells on 64 kPa matrix; p<0.001) (**Fig. 5B**). Treatment of MDA-MB-231 cells with Orai1 inhibitors CM4620 or CM5480 also abrogated the nuclear translocation of YAP (∼50% reduction with CM4620 vs. control, ∼46% reduction with CM5480 vs. control, p<0.001) (**Fig. 5C**), validating the results of genetic KOs. To assess the impact of EHD2-KO or Orai1-KO on YAP/TAZ-TEAD pathway activity, we determined the activity of a transiently transfected TEAD pathway luciferase reporter in cells grown on stiff matrix. Both EHD2-KO and Orai1-KO significantly reduced the reporter activity (∼40% reduction compared to WT, p <0.01; **Fig. 5D**). We further examined the induction of established TEAD target genes, including *CAV1* and *CAVIN1*, the structurally essential components of caveolae, as indicators of the positive feedback loop between caveolae and YAP/TZ-TEAD (Dupont, Morsut et al. 2011, Rausch, Bostrom et al. 2019, Lolo, Walani et al. 2023), using qPCR. The stiff ECM-dependent TEAD target gene expression was significantly impaired by KO of EHD2, or Orai1 (∼52% decrease of *CTGF*, ∼53% decrease of *CYR61*, ∼85% decrease of *ANKRD1*, ∼50% decrease of *CAV1* and ∼40% decrease of *CAVIN1* in EHD2-KO vs. WT; ∼82% decrease of *CTGF*, ∼86% decrease of *CYR61*, ∼90% decrease of *ANKRD1*, 45% decrease of *CAV1* and 40% decrease of *CAVIN1* in Orai1-KO vs. WT; p<0.001) (**Fig. 5E**).

**Figure 5.**
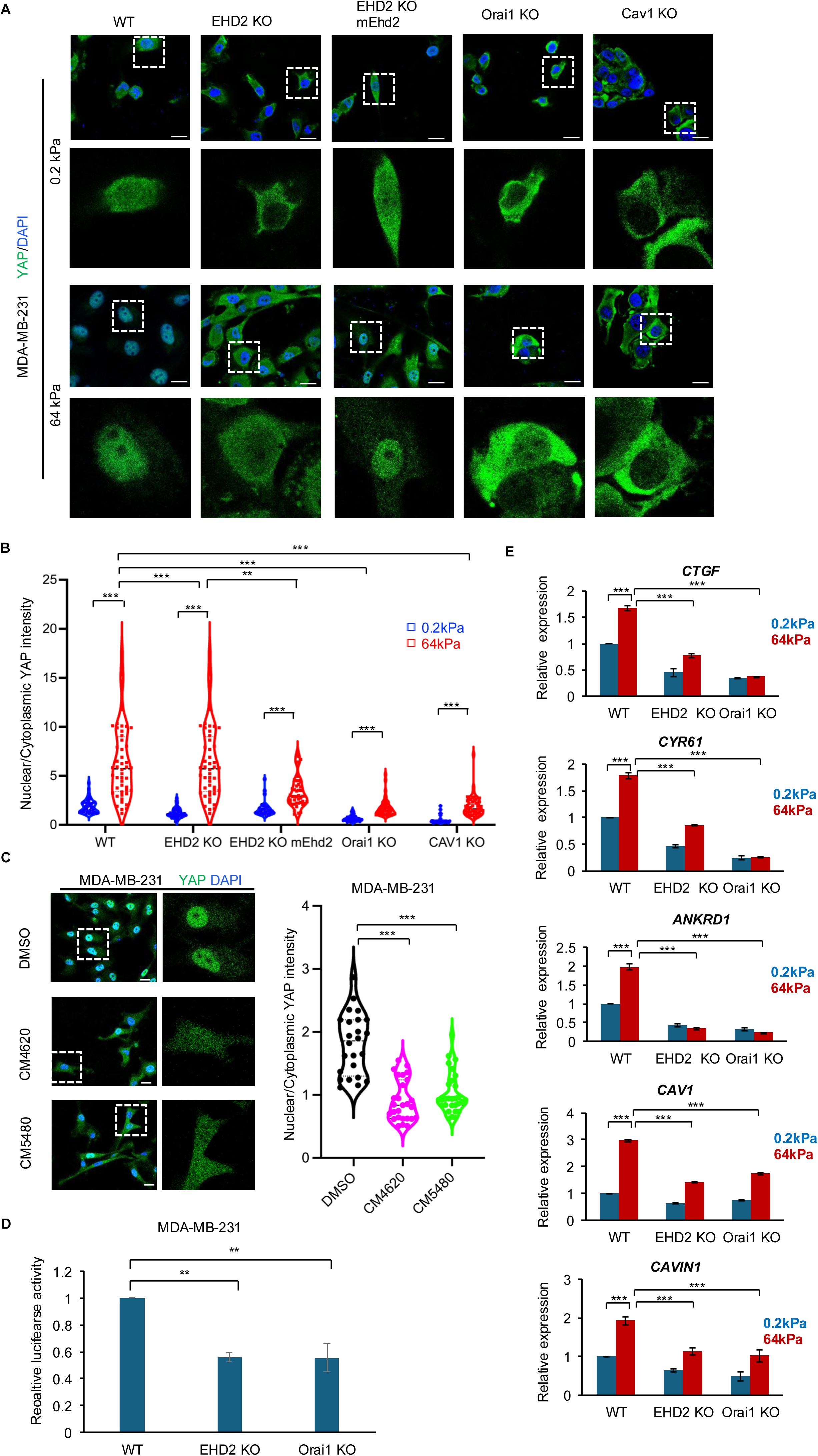
EHD2 and Orai1 are required for mechanosensitive YAP/TAZ pathway activation. **A-B.** Impact of EHD2, Orai1, or CAV1 KO on stiff matrix-induced YAP translocation. The indicated MDA-MB231 wildtype (WT), EHD2-KO, EHD2-KO/mEhd2, Orai1-KO, and CAV1-KO cell lines were cultured on collagen-coated 24-well Cytosoft Rigidity plates layered with 0.2 KPa (soft) or 64 Kpa (stiff) hydrogels, fixed, stained with AF488 conjugated anti-YAP antibody and imaged by confocal imaging. Representative images are shown in **A** (magnified images of a single cell shown under each panel to highlight nuclea/cytoplasmic localization of YAP). Quantification of data is shown in **B**. Image J was used to put masks around the nucleus to quantify fluorescence signals (pixels) within (nuclear) and outside (cytoplasmic) the mask and the data are shown as the ratio of nuclear/cytoplasmic YAP staining signals. Each data point represents cells analyzed in 63 X fields from three independent experiments (at least 39 cells analyzed in each group). One-way ANOVA with Dunnett’s multiple comparisons test, **p<0.01, ***p<0.001. **C**. Impact of Orai1 inhibitors on stiff matrix-induced nuclear localization of YAP. MDAM-MB-231 cells plated on stiff matrix, as in A, were treated for last 4 hours with vehicle (DMSO) or Orai1 inhibitors (CM4620 or CM5480; 10 μM) and cells processed for YAP staining. Nuclear/cytoplasmic ratios of YAP staining were determined as in A/B. Representative images are shown on left. Quantified data of nuclear to cytoplasmic YAP staining ratio are shown on right. Each data point represents cells analyzed from three experiments (n=24). Welch’s t test, ***p<0.001. **D-E**. Impact of EHD2 or Orai1-KO on YAP/TAZ-TEAD pathway gene targets. **D** shows the luciferase activity of transiently transfected YAP/TAZ reporter (pRP-hRluc-8X GTIIC-Luc) in the indicated MDA-MB-231 cell lines cultured on stiff matrix as in A. Data represents mean +/- SEM of three experiments, each with 6 replicates. Welch’s t test, ** p<0.01. **E** shows RT-qPCR analyses of YAP/TAZ downstream genes, *CTGF*, *CYR61*, *ANKRD1*, *CAV1* and *CAVIN1* in the indicated MDA-MB-231 cell lines cultures on 0.2 or 64 Kpa hydrogels. The gene expression values were normalized to GAPDH. Data represents mean +/- SEM of three experiments. One-way ANOVA with Dunnett’s multiple comparisons test, ***p<0.001.

To further confirm the positive feedback between EHD2/Orai1 regulated YAP/TAZ-TEAD pathway activity and the expression of caveolar proteins CAV1 and CAVIN1 seen in TNBC cells with gene KOs, we examined the impact of pharmacological inhibition of TEAD. Treatment of MDA-MB-231 and Hs578T TNBC cells with an allosteric pan-TEAD inhibitor GNE-7883 (Hagenbeek, Zbieg et al. 2023) for 72 hours resulted in a significant reduction in the levels of CAV1 (50% reduction compared to control; p<0.01) (**Fig. 6A**). GNE-7883 treatment in both MDA-MB-231 and Hs578T cell lines significantly reduced the thapsigargin-induced SOCE (28% reduction compared to control in MDA-MB-231 cells and 23% reduction in Hs578T cells; p<0.01) (**Fig. 6B-C**). GNE-7883 treatment of MDA-MB-231 cells for 24 hours also significantly impaired the Ca^2+^ uptake induced by AFM-induced mechanical stimulus to the plasma membrane (6.5-fold Ca^2+^ influx increase in DMSO vs. 2.5-fold Ca^2+^ influx increase in GNE-7883 treatment, p<0.001) (**Fig. 6D**). Finally, short-term GNE-7883 treatment significantly impaired the ability of MDA-MB-231 and Hs578T cells to repair the mechanically-induced (65% repair in control vs. 56% in treated MDA-MB-231 cells at 40 min; 65% repair in control vs. 50% in treated Hs578T cells at 40 min; p <0.05) (**Fig. 6E, Fig. S6**) and SLO-induced (13% PI+ cells in control vs. 17% in treated MDA-MB-231 cells; 35% PI+ cells in control vs. 58% in treated Hs578T cells; p <0.05) (**Fig. 6F**) plasma membrane injuries.

**Figure 6.**
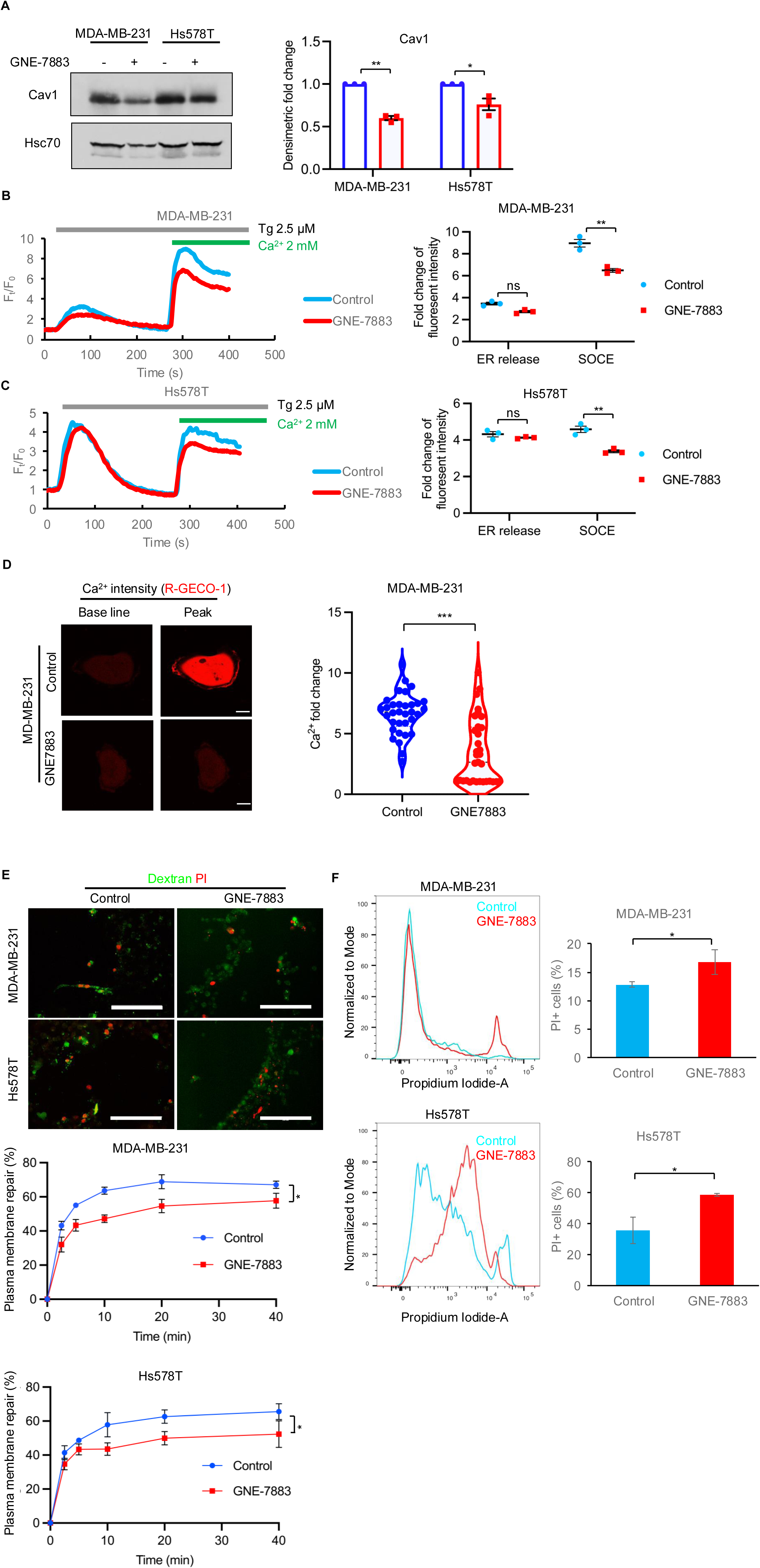
YAP/TAZ-TEAD pathway activity is required for mechanosensitive Ca^2+^ import and membrane repair. **A.** TEAD inhibition reduces CAV1 expression. MDA-MB-231 or Hs578T cells cultured without or with the pan-TEAD inhibitor GNE-7883 (10 μM) for 48 hours were analyzed by WB for CAV1 expression; Hsc70, loading control. Fold-change in signals quantified by densitometry and analyzed using Image J are shown on right. Welch’s t test, *, p<0.05; **, p<0.01. **B-C.** TEAD inhibition reduces the Thapsigargin (TG)-induced SOCE. Cells were pretreated with GNE-7883 (10 μM) for 24 hours. Left, representative plots; right, quantified data from three experiments. Welch’s t test; **p<0.001. **D.** TEAD inhibition impairs the mechanical force-induced Ca^2+^ import. AFM cantilevers were used to apply force and measure Ca^2+^ import from the extracellular space using fluorescence intensity of transfected Ca^2+^ reporter (R-RECO1.2) as the readout. Cells were pretreated with GNE-7883 (10 μM) for 24 hours where indicated. Data points represent cells analyzed through three independent experiments (n=33). Welch’s t test, ***, p<0.001. **E-F.** TEAD inhibition impairs the repair of induced plasma membrane injuries. Shown is the impact of GNE-7883 on the repair of mechanically induced (**E**) **and** SLO-induced (**F**) plasma membrane damage, assessed as in Fig. 1A-B and Fig. 1C-D, respectively. GNE-7883 pre-treatment (10 μM), where indicated, was for 72 hours. Representative confocal images (**E**; top panel) or FACS plots (**F**; right panel) are shown. Quantified data shown are from 3 independent experiments and presented as mean +/- SEM of 3 experiments. Two-way ANOVA or Welch’s t test applied to data in **E** and **F**, respectively, *p<0.05. <u>Scale bar, 20 µm.</u>

Altogether, we find that in addition to an essential role of EHD2-Orai1 axis in acute response to plasma membrane strain, long-term exposure of cells to stiff microenvironment induces EHD2-Orai1 and YAP/TAZ-TEAD axis dependent mechanoadaptation to counteract plasma membrane damage.

## Discussion

The ability to promptly repair plasma membrane breaches is essential for the life of organisms without a cell wall (Cooper and McNeil 2015, Demonbreun and McNally 2016, Andrews and Corrotte 2018). Ca^2+^ inflow from the extracellular space is required to initiate and complete the plasma membrane repair processes (Cheng, Zhang et al. 2015, Cooper and McNeil 2015, Demonbreun and McNally 2016, Andrews and Corrotte 2018, Horn and Jaiswal 2018). The passive inflow of Ca^2+^ from the extracellular space into cytoplasm is thought to be the initial trigger to initiate the repair. Whether plasma membrane-localized Ca^2+^ channels have any role in plasma membrane repair is not known. Studies presented here establish that the plasma membrane Ca^2+^ channel Orai1, a mediator of store-operated Ca^2+^ entry, is essential for mammalian plasma membrane repair. We also establish that EHD2, a caveolae-associated protein required to sustain high plasma membrane levels of Orai1 (Luan, Bielecki et al. 2023), is also essential for mammalian plasma membrane repair. Importantly, we show that EHD2 and Orai1 function as required upstream components of a mechanotransduction cascade to activate YAP/TAZ-TEAD signaling. Collectively, our findings identify novel roles of Orai1-mediated Ca^2+^ transport to sustain Ca^2+^-dependent mammalian plasma membrane repair (Cheng, Zhang et al. 2015, Cooper and McNeil 2015, Demonbreun and McNally 2016, Andrews and Corrotte 2018, Horn and Jaiswal 2018) and for activation of YAP/TAZ-TEAD pathway of mechanoadaptation (Dupont, Morsut et al. 2011, Rausch, Bostrom et al. 2019, Lolo, Walani et al. 2023).

The role of Orai1 (or its family members Orai2/3), in partnership with STIM1 (or STIM2) in SOCE is well-established, but this role has almost exclusively been investigated in the context of responses to biochemical signaling, such as cell surface receptor activation (Ong, Subedi et al. 2019, Lewis 2020). In contrast, there is little evidence for a primary role for Orai channels in cellular responses to mechanical stimuli. However, prior studies have suggested the role of Orai1 secondary to activation of other mechanosensitive Ca^2+^ channels. For example, Ca^2+^ entry induced by mechanosensitive Piezo1 channel agonist Yoda-2 in endometrial mesenchymal stem cells was shown to be dampened by 2-APB, an inhibitor of Orai channel-mediated SOCE (Chubinskiy-Nadezhdin, Semenova et al. 2022). In a study using an in vivo paradigm of flow shear-dependent mechanotransduction, Orai1-KO embryos displayed reduced lymphatic density and impaired lymphatic development (Choi, Park et al. 2017). Further studies demonstrated that Piezo1 channel served in the mechanosensory role, with Orai1 mediating the subsequent SOCE (Choi, Park et al. 2017). We show that direct application of mechanical force to the plasma membrane initiates rapid Ca^2+^ import that is completely abrogated by Orai1-KO as well as by Orai1 channel-selective inhibitors (**Fig. 4A-C**). We have previously established that loss of EHD2 leads to reduced SOCE, with a reduction in the plasma membrane pool of Orai1 while the total Orai1 levels were unchanged (Luan, Bielecki et al. 2023). Indeed, the mechanosensitive Ca^2+^ entry was also lost upon EHD2-KO (**Fig. 4B**). Consistent with Orai1 localization to CAV1-containing plasma membrane domains (Sathish, Abcejo et al. 2012, Chantôme, Potier-Cartereau et al. 2013, Jardin and Rosado 2016, Bohorquez-Hernandez, Gratton et al. 2017), we observed loss of mechanosensitive Ca^2+^ entry upon CAV1-KO (**Fig. 4B**). Mechanosensitivity of Ca^2+^ import through G-GECO1-Orai1, a genetically-encoded Ca^2+^ biosensor that reports direct Ca^2+^ influx through Orai1 (Dynes, Amcheslavsky et al. 2016), (**Fig. 4D**) further substantiates the role of Orai1 in mechanosensitive Ca^2+^ entry across the plasma membrane. We further demonstrate the mechano-sensitive, Orai1-mediated Ca^2+^ entry to require STIM proteins, indicating that such Ca^2+^ transport occurs at ER-PM contacts, where STIM proteins are known to interact with and activate Orai1 (Ong, Subedi et al. 2019, Lewis 2020). Collectively, our results define a new mechanosensitive role for plasma membrane caveolae-localized Orai1. The absence of a retained intracellular store Ca^2+^ release component upon mechanical indentation of Orai1-, EHD2- or CAV1-KO cell models argues against the likelihood of a distinct mechanosensitive Ca^2+^ channel mediating the initial Ca^2+^ release with Orai1 functioning in a classical role as an SOCE channel in our cell system. However, given the findings in Orai1-KO mice discussed above, it remains possible that mechanosensitive proteins, including the known mechanosensitive Ca^2+^ channels, are involved in Orai1 activation in response to mechanical stimuli. A systematic analysis of candidate mechanosensitive Ca^2+^ channels and accessory proteins will be needed to clarify this further.

Given the accumulation of EHD2 at plasma membrane repair sites in skeletal muscle models (Marg, Schoewel et al. 2012, Demonbreun, Quattrocelli et al. 2016), our findings in cancer cells suggest a comparable role of EHD2 in other cell types with known roles of caveolae in plasma membrane repair (Sinha, Koster et al. 2011, Corrotte, Fernandes et al. 2012, Corrotte, Almeida et al. 2013, Shvets, Ludwig et al. 2014, Cheng, Mendoza-Topaz et al. 2015, Garcia, Bagwell et al. 2017). More importantly, our results raise the possibility of a broader and essential role of Orai1 in caveolae-dependent repair across cell types and organisms. Whether EHD2 functions as an obligate partner in such a role remains to be determined; while EHD2 and CAV1 proteins are coordinately expressed in breast cancer cell lines (Luan, Bielecki et al. 2023) and *CAV1/CAVIN1* and *EHD2* mRNAs show strong co-expression across cell types, *EHD2* and *Orai1* mRNA expression is not similarly correlated (based on single cell portal and mRNA co-expression databases). Consistent with EHD2-independent Orai1 function in membrane repair, the residual repair in EHD2-KO cells was still Ca^2+^ dependent (**Fig. 1B**), likely reflecting the reduction but not a complete absence of cell surface Orai1 in these cells (Luan, Bielecki et al. 2023). Further analyses in naturally EHD2-low/non-expressing cell systems will be required to explore this further.

Prior work has established the YAP/TAZ-TEAD pathway as a major mechanosensitive signaling axis in response to ECM stiffness, shear stress and cell stretching in a manner independent of the upstream Hippo pathway kinases (Dupont, Morsut et al. 2011, Rausch, Bostrom et al. 2019, Lolo, Walani et al. 2023). CAV1 was identified as a critical upstream positive regulator of such mechanosensitive YAP/TAZ-TEAD pathway activation (Moreno-Vicente, Pavon et al. 2018). Consistent with the required role of EHD2 to stabilize plasma membrane caveolae (Sathish, Abcejo et al. 2012, Chantôme, Potier-Cartereau et al. 2013, Jardin and Rosado 2016, Bohorquez-Hernandez, Gratton et al. 2017), we found that deletion of EHD2 impaired the stiff ECM dependent YAP nuclear translocation, phenocopying the impact of CAV1-KO (**Fig. 5A-B**). Importantly, we found that Orai1-KO or its inhibition also impaired the YAP translocation induced by stiff ECM (**Fig. 5C**). Further, EHD2-KO or Orai1-KO reduced the stiff ECM induced TEAD target gene expression (**Fig. 5D**). These findings establish a novel requirement of EHD2 and Orai1 in mechanosensitive YAP/TAZ-TEAD activation. Combining insights from our earlier study linking EHD2 and Orai1 (Luan, Bielecki et al. 2023), and our findings here, we propose that EHD2-mediated stabilization of CAV1-containing plasma membrane domains places Orai1 at mechanosensitive plasma membrane domains and that mechanical stimuli activate Orai1 as a required step in YAP/TAZ-TEAD pathway activation. Orai1-mediated Ca^2+^ import in the context of cell surface receptor activation as a trigger to regulate multiple cellular signaling pathways is well established, including several transcriptional regulatory pathways (Nieto-Felipe, Macias-Diaz et al. 2023). The critical role of Orai1 as an upstream positive regulator of YAP/TAZ-TEAD pathway activation, as we identify here, therefore provides a novel paradigm to understand CAV1-dependent mechanosensitive signaling. Consistent with this suggestion, mechanosensitive YAP/TAZ-TEAD activation was found to require Rho GTPase activity and actomyosin cytoskeletal contractility (Dupont, Morsut et al. 2011), which in turn are known be regulated by mechanosensitive Ca^2+^ fluxes (Higashida, Kiuchi et al. 2013, Pardo-Pastor, Rubio-Moscardo et al. 2018, Lakk and Krizaj 2021, Miroshnikova, Manet et al. 2021, Varadarajan, Chumki et al. 2022, Fu, Wang et al. 2024).

Prior studies have shown that YAP/TAZ-TEAD pathway is required for the expression of structural components of caveolae, CAV1 and CAVIN1, and inhibition of YAP/TAZ-TEAD axis led to loss of plasma membrane caveolae (Dupont, Morsut et al. 2011, Rausch, Bostrom et al. 2019, Lolo, Walani et al. 2023). More recently, mild to moderate mechanical plasma membrane stress was shown to activate YAP/TAZ-TEAD pathway through CAV1-containing but CAVIN1-negative plasma membrane dolines as a mechanoadaptation mechanism through feedback upregulation of CAV1 and CAVIN1 gene expression to increase the CAV1/CAVIN1-containing plasma membrane caveolae, which are required for mechanical protection against more severe plasma membrane stress (Dupont, Morsut et al. 2011, Rausch, Bostrom et al. 2019, Lolo, Walani et al. 2023). Consistent with EHD2-Orai1 axis as an intermediary in such mechanoadaptive positive feedback, stiff matrix-induced CAV1 and CAVIN1 gene expression was reduced by EHD2 or Orai1 KO (**Fig. 5E**). Further supporting this mechanism downstream of the EHD2-Orai1 axis, even short-term pharmacological TEAD inhibition reduced the levels of CAV1/CAVIN1 proteins (**Fig. 6A**) and the SOCE response elicited upon ER Ca^2+^ store depletion (using thapsigargin) (**Fig. 6B-C**), Remarkably, TEAD inhibition significantly impaired the Ca^2+^ import in response to plasma membrane force application (**Fig. 6E-F**), expanding the YAP/TAZ-TEAD mediated feedback to the EHD2-Orai1-dependent mechanosensitive Ca^2+^ import, which in turn our findings establish as essential for YAP/TAZ-TEAD pathway activation.

Our findings that EHD2 is required for rapid Orai1 activation in response to mechanical force applied to the plasma membrane (**Fig. 4A**) are consistent with the requirement of EHD2 to sustain the plasma membrane pool of caveolae (Sathish, Abcejo et al. 2012, Chantôme, Potier-Cartereau et al. 2013, Jardin and Rosado 2016, Bohorquez-Hernandez, Gratton et al. 2017) and our previous work that EHD2 is required to sustain high plasma membrane levels of Orai1 (Luan, Bielecki et al. 2023). However, our finding of rapid EHD2 recruitment to plasma membrane sites of applied mechanical force (**Fig. 3B**) differs from previous findings of CAV1-dependent modest or substantial release of EHD2 from the plasma membrane in response to cyclical stretch or hypotonic stress, respectively, with sumo modification of EHD2 leading to its nuclear localization (Torrino, Shen et al. 2018). The discordant results may reflect our use of transiently applied localized mechanical force as opposed to more prolonged cell-wide mechanical stimulation in prior studies and will need further investigation. That mechanosensitive Orai1 activation, which occurs at the plasma membrane, requires EHD2, strongly argues for the observed mechanosensitive role of EHD2 at the plasma membrane rather than through nuclear localization. In prior work, we found that high nuclear staining of EHD2 in breast cancer tissues was associated with longer patient survival, diametrically opposite to the association of high non-nuclear EHD2 overexpression with shorter patient survival (Luan, Bielecki et al. 2023). Thus, the nuclear localization of EHD2 in response to mechanical stress may reflect its sequestration for later utilization in plasma membrane-associated functions. Indeed, the prior study discussed above (Torrino, Shen et al. 2018) found rapid exit of EHD2 from the nucleus and its re-localization to plasma membrane during recovery from hypotonic stress. Further, NIH-3T3 cells rendered caveolae-deficient by the combined EHD1, 2 and 4 KO exhibited susceptibility or membrane ruptures upon prolonged cyclical stretching (Yeow, Howard et al. 2017).

An unresolved question is how the mechanical force applied to plasma membrane might activate Orai1 in an EHD2-dependent and STIM-dependent manner. Changes in membrane curvature sensed by curvature sensing proteins are well known to affect cellular responses (McMahon and Boucrot 2015). For example, mechanosensitive Piezo1 Ca^2+^ channel was found enriched at plasma membrane invaginations and depleted at filopodia (Yang, Miao et al. 2022). Recently, use of vertical pillars to induce plasma membrane curvature changes mimicking the cardiomyocyte plasma membrane transverse tubules, sites enriched for ER-PM contacts, was found to induce ER-PM contacts through junctophilin proteins; interaction of junctophilin-2 with EHD proteins was identified as a mechanism for curvature sensing to promote ER-PM contact enrichment (Yang, Valencia et al. 2024). Thus, it is plausible that mechanical force induced curvature on the plasma membrane recruits EHD2 to promote rapid ER-PM contact formation or stabilization to promote STIM1-Orai1 interaction and mechanosensitive Ca^2+^ entry. While studies in cardiomyocytes showed junctophilin-2 interaction with multiple EHD proteins, and EHD4 was functionally implicated, these other family members are expressed in parental as well as EHD2-KO breast cancer cell models used here (Luan, Bielecki et al. 2023) and do not appear to compensate for the role of EHD2. It is possible that the requirement of EHD2 for high PM Orai1 expression (Luan, Bielecki et al. 2023) contributes to this relative specificity.

In conclusion, our findings establish the EHD2-Orai1 duo as a novel regulator of mechanosensitive import of extracellular Ca^2+^ essential for efficient mammalian cell plasma membrane repair. We also establish the EHD2-Orai11 axis as a critical upstream element required for the activation of mechanosensitive YAP/TAZ-TEAD dependent gene expression for mechanoadaptation. Further studies of this novel plasma membrane mechanosensory apparatus are likely to reveal key new insights into mechanotransduction and regulation of plasma membrane homeostasis under physiological states and in diseases, such as cancer.

## ACKNOWLEDGEMENTS

We thank the Band Lab members for their many discussions, and the staff of UNMC Core facilities for their invaluable assistance. We thank Drs. Robert Campbell (CMV-R-GECO1.2), Michael Cahalan (G-GECO1-Orai1) and Ari Helenius (EHD2-mCherry) for plasmids obtained through Addgene. This work was supported by grants from Department of Defense breast cancer research program (W81XWH-17-1-0616 and W81XWH-20-1-0058 to HB, and W81XWH-20-1-0546, HT94252410337 and HT9425-23-1-0052 to VB), the NIH (R21CA297629 to HB, R21CA241055 and R03CA253193 to VB, and P20 GM121316 to SPR), a UNMC startup grant (to SPR), the Fred & Pamela Buffett Cancer Center pilot grants (to HB & VB), a Nebraska Research Initiative seed grant (to HB and JYL), a Nebraska DHHS LB506 pilot award (18123-Y3 to HB), a Children’s Hospital Research Institute pilot grant (to HB), and the Raphael Bonita Memorial Fund (to HB). Research reported in this publication was supported by the Advanced Microscopy, Flow Cytometry, and Data Science Shared Core Facilities which are partially funded by the National Cancer Institute of the National Institutes of Health Cancer Center Support Grant to the Fred & Pamela Buffett Cancer Center (P30CA036727) and the Nebraska Research Initiative. TAB was a trainee under the National Cancer Institute Cancer Biology Training Grant T32CA009476.

## Author Contributions

HL (Investigation, data curation, formal analysis, methodology, visualization, writing – original draft); AC, TAB, BCM, SS, YW, MDS (Investigation, data curation, formal analysis, methodology); LMS (Formal analysis); KAS (Resources); DWC & AK (Funding acquisition), JD (Resources, methodology), JTL (Conceptualization, funding acquisition, writing – review and editing), VB (Conceptualization, funding acquisition, supervision, writing – original draft, review & editing), SPR (Conceptualization, supervision, investigation, formal analysis, visualization, writing – original draft; review & editing), HB (Conceptualization, funding acquisition, supervision, resources, formal analysis, writing – original draft, review & editing).

## DECLARATION OF INTERESTS

KAS is an employee of CalciMedica Inc., which is advancing Orai1 inhibitors into clinic. Other authors have nothing to declare.

### Materials availability

All unique/stable reagents generated in this study are available from the lead contact with a completed materials transfer agreement. The recipient will incur shipping costs.

## STAR METHODS

### Key resources table

| REAGENT OR RESOURCE | SOURCE | IDENTIFIER |
| --- | --- | --- |
| Antibodies |  |  |
| Orai1 | Millipore-Sigma | Cat# O8264; (Luan, Bielecki et al. 2023) |
| Beta-actin | Millipore-Sigma | Cat# SAB11305567 |
| Hsc70 | Santa Cruz | Cat# sc-7298 |
| Caveolin1 | BD Biosciences | Cat# 610057 |
| EHD2 | (Luan, Bielecki et al. 2023) | N/A |
| YAP (IF) | Cell Signaling Technology | Cat# 14729 |
| HRP-conjugated Protein A antibody | ThermoFisher Scientific | Cat# 101023 |
| HRP-conjugated goat anti-mouse antibody | ThermoFisher Scientific | Cat# 31431 |
| Chemicals, peptides, and recombinant proteins |  |  |
| Thapsigargin | ThermoFisher Scientific | Cat# T7459 |
| Fluo 4 AM | ThermoFisher Scientific | Cat# 14201 |
| CM4620 | SelleckChem | Cat# S6834 |
| CM5480 | CalciMedica | N/A |
| GsMTx4 | SelleckChem | Cat# P1205 |
| Yoda2 | TOCRIS | Cat# 8051 |
| TRIZol | ThermoFisher Scientific | Cat# 15596026 |
| Critical commercial assays |  |  |
| Dual-Luciferase® Reporter assay Kit | Promega | Cat# E1910 |
| RT-qPCR kit | Qiagen | Cat# 204141 |
| Experimental models: Cell lines |  |  |
| MDA-MB-231 | ATCC | Cat# HTB-26 |
| Hs578T | ATCC | Cat# HTB-126 |
| MDA-MB-231 EHD2 KO | (Luan, Bielecki et al. 2023) | N/A |
| Hs578T EHD2 KO | (Luan, Bielecki et al. 2023) | N/A |
| MDA-MB-231 CAV1 KO | (Luan, Bielecki et al. 2023) | N/A |
| MDA-MB-231 EHD2 KO *mEhd2* | (Luan, Bielecki et al. 2023) | N/A |
| Recombinant DNA |  |  |
| CMV-R-GECO1.2 | (Wu, Liu et al. 2013) | Addgene# 45494 |
| G-GECO1-Orai1 | Joseph L Dynes et al. 2015 (Dynes, Amcheslavsky et al. 2016) | Addgene# 73561 |
| EHD2-mChery | GeneCopoeia | Cat# EX-A3485-Lv155 |
| pLenti-U6-sgRNA-SFFV-Cas9-2A-Puro (Orai1 KO generation) | Abm | Cat# 35720125 |
| Software and algorithms |  |  |
| ImageJ | https://imagej.net/software/imagej/index | N/A |
| FlowJo 10 | FlowJo | N/A |
| Zen | Zeiss | N/A |
| Prism 9 | GraphPad | N/A |
| Oligonucleotides |  |  |
| See Table S1 for RT-qPCR primers | N/A | N/A |

## EXPERIMENTAL MODEL AND SUBJECT DETAILS

### Cell lines and medium

MDA-MB-231 cell line (obtained from ATCC) was cultured in complete α-MEM medium with 5% fetal bovine serum, 10 mM HEPES, 1 mM each of sodium pyruvate, nonessential amino acids, and L-glutamine, 50 μM 2-ME, and 1% penicillin/ streptomycin (Life Technologies, Carlsbad, CA). Hs578T cell line (ATCC) was cultured in α-MEM medium supplemented as above plus 1 μg/mL hydrocortisone and 12.5 ng/mL epidermal growth factor (Millipore Sigma, St. Louis, MO). Generation and maintenance of EHD2-KO TNBC cell lines, EHD2-KO cell lines reconstituted with mouse EHD2 (EHD2-KO-mEHD2) and CAV1-KO cell lines have been described previously (Luan, Bielecki et al. 2023).

### Antibodies and reagents

Antibodies used for immunoblotting were as follows: Orai1 (# O8264) and beta-actin (# SAB1305567) from Millipore-Sigma; HSC70 (# sc-7298) from Santa Cruz Biotechnology; Caveolain-1 (#610057) from BD Biosciences; Cavin 1(#46379) from Cell Signaling Technology. Horseradish peroxidase (HRP)-conjugated Protein A or HRP-conjugated goat anti-mouse secondary antibody for immunoblotting were from Invitrogen. YAP antibody (Alexa Fluor® 488 Conjugate, #14729) for immunofluorescence (IF) staining was from Cell Signaling Technology. Thapsigargin (# T7459) and Fluo 4AM (#14201) were from ThermoFisher Scientific. Orai1 inhibitor CM4620 (Waldron, Chen et al. 2019) was from SelleckChem (#S6834); Orai1 inhibitor CM5480 (Szabo, Csakany-Papp et al. 2023) was provided by CalciMedica Inc. (La Jolla, CA).

### Transfection reagents and plasmids

XtremeGENE 9 transfection reagent was from Roche Applied Science (Indianapolis, IN); CMV-R-GECO1.2 (Wu, Liu et al. 2013) was a gift from Robert Campbell (Addgene plasmid # 45494; http://n2t.net/addgene:45494; RRID:Addgene_45494). G-GECO1-Orai1 (Dynes, Amcheslavsky et al. 2016) was a gift from Michael Cahalan (Addgene plasmid # 73561; http://n2t.net/addgene:73561; RRID:Addgene_73561). EHD2-mChery plasmid (# EX-A3485-Lv155) was from GeneCopoeia.

### Generation of CRISPR-Cas9 knockout cell lines

All-in-One sgRNA CRISPR/Cas9 Lentivectors from Applied Biological Materials (Richmond, BC, Canada) were used to derive Orai1 (pLenti-U6-sgRNA-SFFV-Cas9-2A-Puro, #35720125) KO cell lines.

### SOCE assay

Cells were seeded in 35 mm glass-bottom dishes (cat. #FD35-100, WPI Inc) and loaded with Fluo4-AM in modified Tyrode’s solution (2 mM calcium chloride, 1 mM magnesium chloride, 137 mM sodium chloride, 2.7 mM potassium chloride, 12 mM sodium bicarbonate, 0.2 mM sodium dihydrogen phosphate, 5.5 mM glucose, pH 7.4) for 1 hour at 37oC. After washing with Ca^2+^-free Tyrode’s solution, live cells were imaged under a confocal microscope (LSM710; Carl Zeiss), with fluorescence excitation at 488 nm and emission at 490–540 nm. To initiate Ca^2+^ release from intracellular stores, 2.5 μM thapsigargin was added in the absence of extracellular Ca^2+^. Once the Ca^2+^ signals approached the baseline, calcium chloride was added to 2 mM final concentration to record the SOCE. Data is presented as fold change in fluorescence emission relative to baseline.

### Membrane repair assay

For mechanical injury, confluent cell monolayers in 48-well plates were incubated in Tyrode’s solution containing membrane-impermeable FITC-Dextran (500 µg/mL). A standardized mechanical injury was introduced via a single scratch wound using a 200 µL pipette tip, allowing dye entry into cells with compromised plasma membranes. At designated time points post-injury, the extracellular dye was removed followed by a PBS wash. Cells were then briefly incubated with propidium iodide (PI; 50 µg/mL) to label the nuclei of cells with unrepaired plasma membrane defects. Imaging was performed via fluorescence microscopy. FITC+ cells represented the total population of injured cells. FITC+PI+ (double-positive) cells were scored as those with failure to repair their membrane defects. The repair efficiency was calculated as the percentage of injured (FITC+) cells that excluded PI (FITC+PI-) at each time point. For Streptolysin O (SLO) induced membrane injury, 106 trypsin/EDTA-released and washed cells were incubated with SLO (25 U/mL) in suspension for 5 min at 4°C in 250 μl of Ca^2+^-free Tyrode’s solution followed by resuspension in 37°C Tyrode’s solution for 10 min and PI staining. After flow cytometry (FACSCalibur; Becton Dickinson) of at least 10,000 cells, the data were analyzed using the FlowJo software (Tree Star, Inc.).

### Plasma membrane indentation using Atomic Force Microscopy (AFM)

Cells were cultured on 35 mm glass bottom dishes and transfected with R-GECO1.2 (Wu, Liu et al. 2013) or EHD2-mCherry (Stoeber, Stoeck et al. 2012) plasmid. Live cell imaging was conducted with cells were in CO₂-independent medium supplemented with 10% FBS and 1% penicillin–streptomycin and maintained at 37 °C with a JPK Petri Dish Heater to preserve the physiological conditions. Indentation was carried out on a Bruker CellHesion 200 AFM using a pyramidal-tipped microcantilever (FM-10, NanoAndMore). A loading rate of 3 µm/s and a maximum contact force of 100 nN were applied, with each indentation held for 20 s. Simultaneous high-resolution imaging was performed using a Zeiss LSM 900 confocal microscope. AFM force–distance data were processed in JPK Data Processing (JPK-DP), while confocal images were analyzed using FIJI/ImageJ. Statistical analyses and plotting were conducted in OriginLab Pro.

### Immunofluorescence microscopy

Cells were cultured on collagen (#5005, Advanced Biomatrix) pre-coated glass bottom CytoSoft® Imaging 24-Well Plate of different stiffness (#5183 for 0.2 kPa, #5189 for 64k Pa, Advanced Biomatrix) to about 50% confluency, fixed with 4% PFA/PBS (10 min), blocked with 5% BSA/PBS (60 min), and incubated with Fluorescent conjugated antibodies in 5% BSA/PBS overnight at 4 °C. Nuclei were visualized with Hoechst 33342 (#62249, ThermoFisher Scientific) staining. Fluorescence images were captured on a Zeiss LSM-800 confocal microscope (63X objective) and analyzed using the ZEN software (Zeiss).

### Western blotting

Cells were lysed in Triton-X-100 lysis buffer (50 mM Tris pH 7.5, 150 mM NaCl, 0.5% Triton-X-100, 1 mM PMSF, 10 mM NaF, and 1 mM sodium orthovanadate). Lysates were rocked at 4 °C for at least 1 hr, spun in a microfuge at 13,000 rpm for 20 min at 4°C and supernatant protein concentration determined using the BCA assay kit (Thermo Fisher Scientific, Rockford, IL). 50 μg aliquots of lysate proteins were resolved on sodium dodecyl sulfate-7.5% or 12% polyacrylamide gel electrophoresis (SDS-PAGE), transferred to polyvinylidene fluoride (PVDF) membrane, and immunoblotted with the indicated antibodies.

### Trans-well migration assay

Cells grown in 0.5% FBS-containing starvation medium for 24 h were trypsinized and seeded at 10^4^ on top chambers of 24-well plate trans-wells (# 353097, Corning) in 200 μL of growth factor deprived medium. After 3 h, medium containing 10% FBS was added to lower chambers and trans-wells incubated at 37oC for 16 h. The non-migrated cells on the upper surface of membranes were removed with cotton swabs, and the migrated cells on the lower surface methanol-fixed and stained in 0.5% crystal violet in methanol. Six random 10× fields per insert were photographed, and cells counted using the ImageJ software. Each experiment was run in triplicates and repeated three times.

### YAP/TAZ-TEAD pathway luciferase reporter assay

Cells were transfected with a synthetic TEAD dual luciferase reporter (pRP-hRluc-8X GTIIC-Luc, cat# VB250204-1311 from VectorBuilder). All luciferase emission measurements were performed using a Dual-Luciferase® Reporter assay Kit (DLR™ assay, Promega). Luminescence was recorded using a GloMax® luminometer (Promega).

### Quantitative real-time PCR

Total RNA was extracted using TRIzol reagent (#15596026, Invitrogen), reverse transcribed using a real-time Quantitative PCR kit (#204141, Qiagen) and used for real-time QPCR with primers listed in **Table 1**.

**Table 1.**
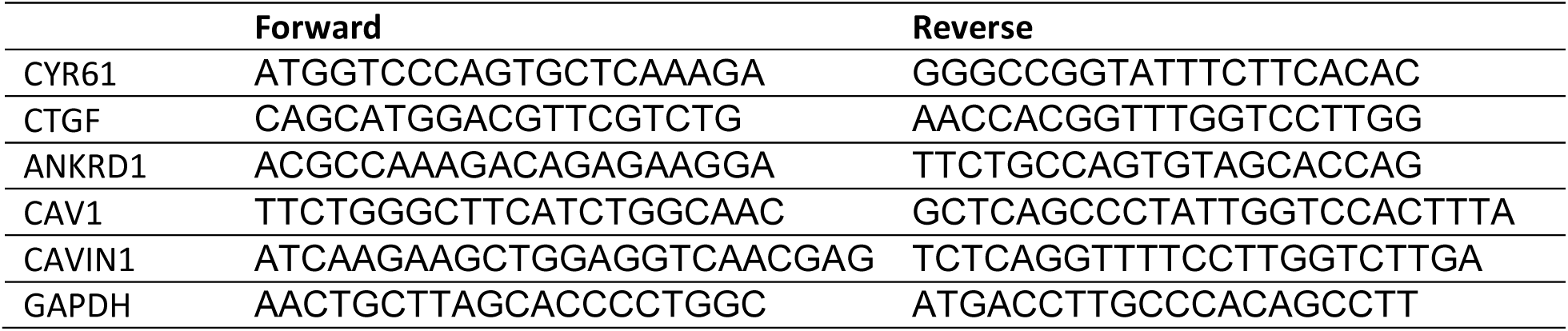
Primer sequences for RT-qPCR.

### Statistical analysis

GraphPad Prism software (version 9) was used to perform statistical analyses. Statistical analysis of cell biological data was performed by comparing groups using unpaired Welch’s *t*-test (two groups), one-way ANOVA with Dunnett’s multiple comparisons test (more than two groups) and two-way ANOVA test (two factors). *p* values of <0.05 were considered significant.

---

**Fig S1.**
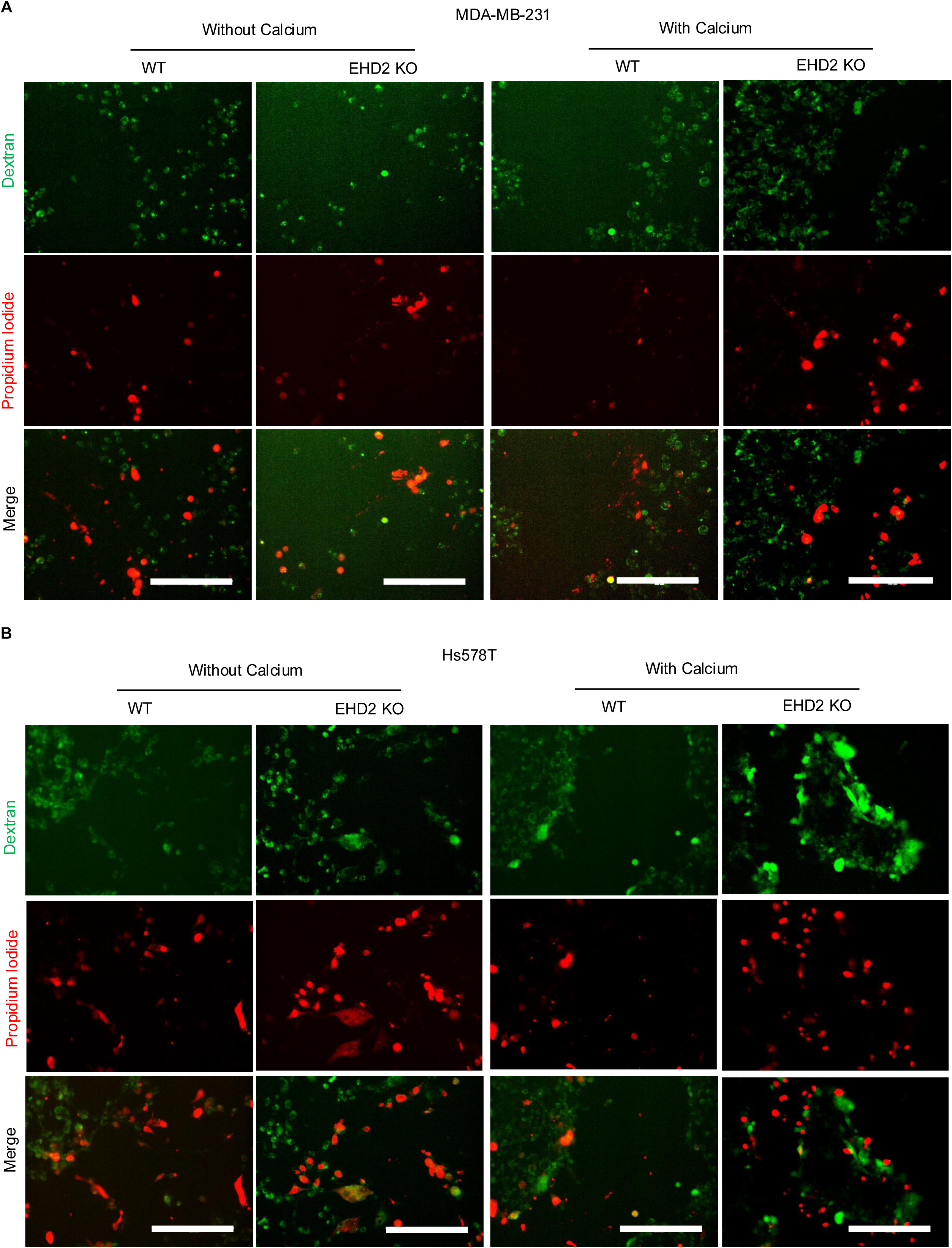

**Fig S2.**
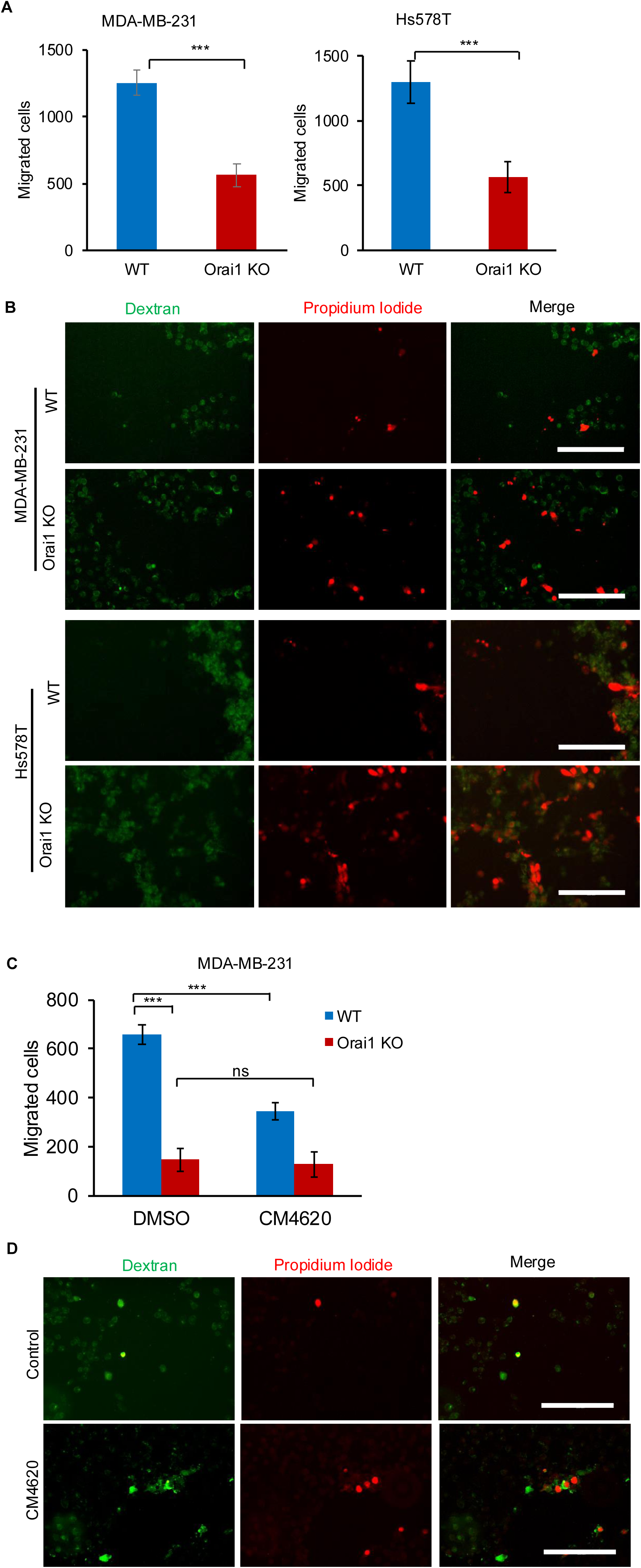

**Fig S3.**
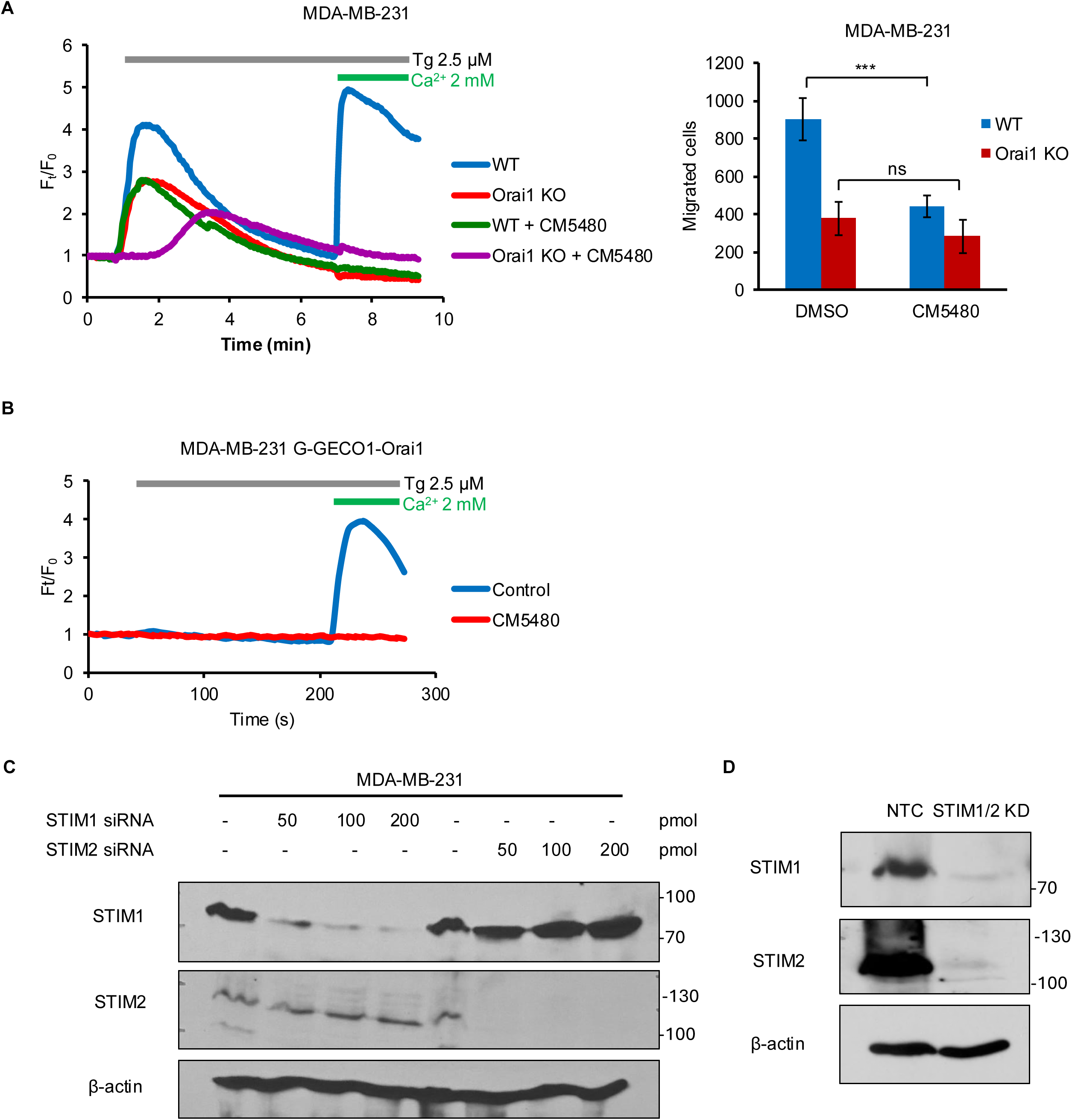

**Fig S4.**
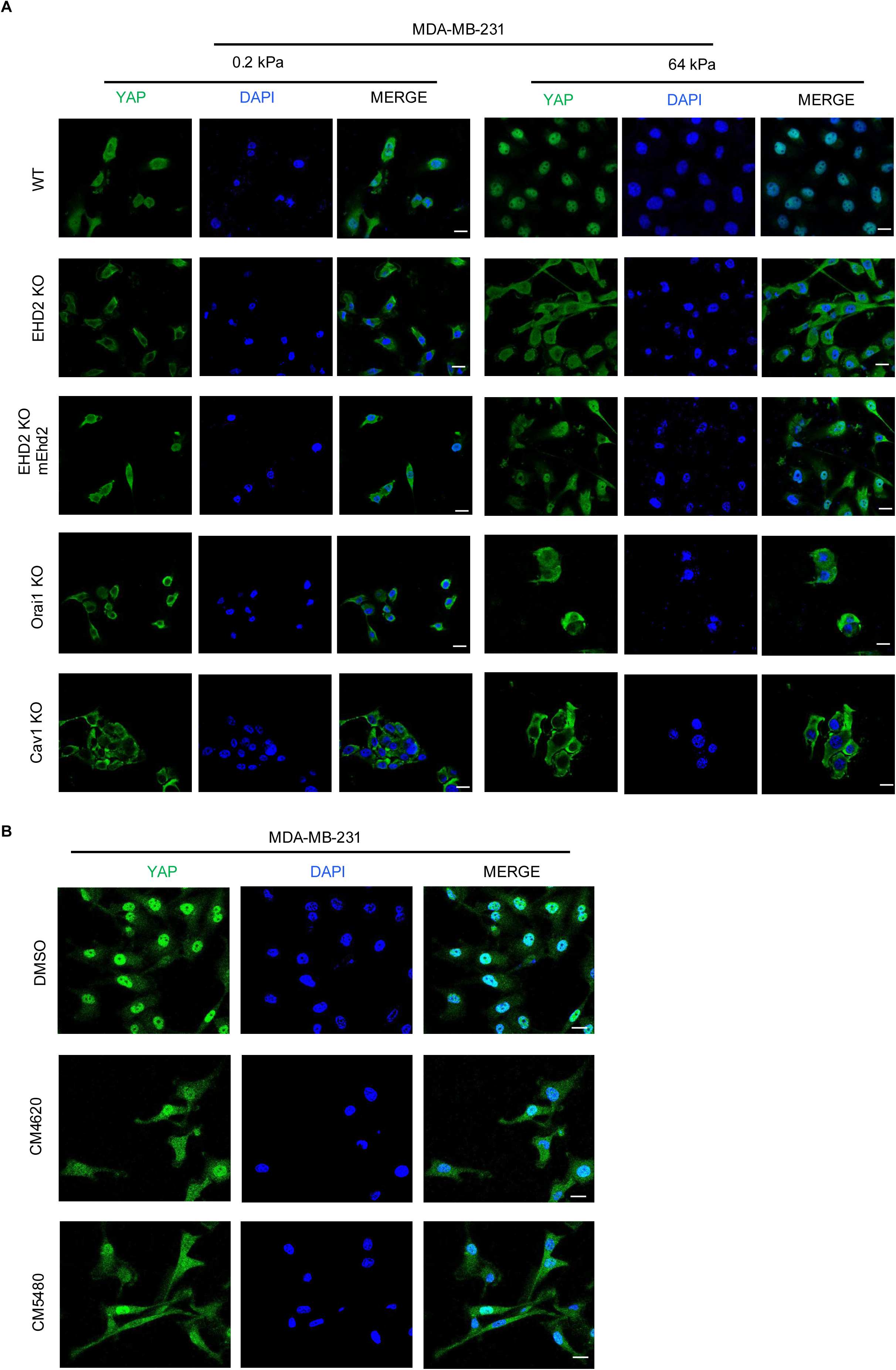

**Fig S5.**
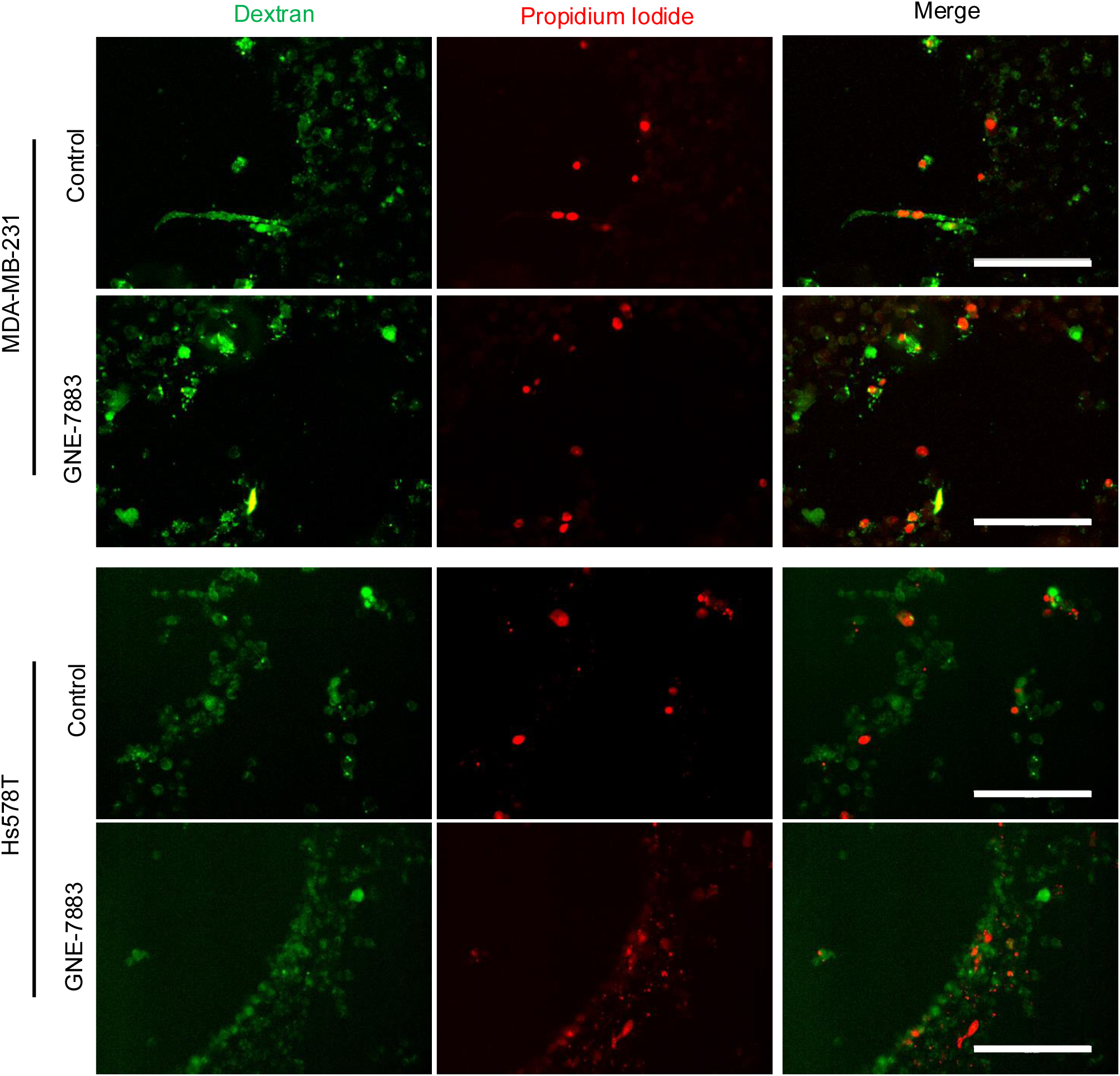

## References

Andrews, N. W. and M. Corrotte (2018). “Plasma membrane repair.” Curr Biol 28(8): R392–R397.

Bohorquez-Hernandez, A., E. Gratton, J. Pacheco, A. Asanov and L. Vaca (2017). “Cholesterol modulates the cellular localization of Orai1 channels and its disposition among membrane domains.” Biochim Biophys Acta Mol Cell Biol Lipids 1862(12): 1481–1490.

Bouvet, F., M. Ros, E. Bonedeau, C. Croissant, L. Frelin, F. Saltel, V. Moreau and A. Bouter (2020). “Defective membrane repair machinery impairs survival of invasive cancer cells.” Sci Rep 10(1): 21821.

Bruen, C., M. Al-Saadi, E. A. Michelson, M. Tanios, R. Mendoza-Ayala, J. Miller, J. Zhang, K. Stauderman, S. Hebbar and P. C. Hou (2022). “Auxora vs. placebo for the treatment of patients with severe COVID-19 pneumonia: a randomized-controlled clinical trial.” Crit Care 26(1): 101.

Bruen, C., J. Miller, J. Wilburn, C. Mackey, T. L. Bollen, K. Stauderman and S. Hebbar (2021). “Auxora for the Treatment of Patients With Acute Pancreatitis and Accompanying Systemic Inflammatory Response Syndrome: Clinical Development of a Calcium Release-Activated Calcium Channel Inhibitor.” Pancreas 50(4): 537–543.

Chantôme, A., M. Potier-Cartereau, L. Clarysse, G. Fromont, S. Marionneau-Lambot, M. Guéguinou, J. C. Pagès, C. Collin, T. Oullier, A. Girault, F. Arbion, J. P. Haelters, P. A. Jaffrès, M. Pinault, P. Besson, V. Joulin, P. Bougnoux and C. Vandier (2013). “Pivotal role of the lipid Raft SK3-Orai1 complex in human cancer cell migration and bone metastases.” Cancer Res 73(15): 4852–4861.

Cheng, J. P., C. Mendoza-Topaz, G. Howard, J. Chadwick, E. Shvets, A. S. Cowburn, B. J. Dunmore, A. Crosby, N. W. Morrell and B. J. Nichols (2015). “Caveolae protect endothelial cells from membrane rupture during increased cardiac output.” J Cell Biol 211(1): 53–61.

Cheng, X., X. Zhang, Q. Gao, M. Ali Samie, M. Azar, W. L. Tsang, L. Dong, N. Sahoo, X. Li, Y. Zhuo, A. G. Garrity, X. Wang, M. Ferrer, J. Dowling, L. Xu, R. Han and H. Xu (2014). “The intracellular Ca(2)(+) channel MCOLN1 is required for sarcolemma repair to prevent muscular dystrophy.” Nat Med 20(10): 1187–1192.

Cheng, X., X. Zhang, L. Yu and H. Xu (2015). “Calcium signaling in membrane repair.” Semin Cell Dev Biol 45: 24–31.

Choi, D., E. Park, E. Jung, Y. J. Seong, M. Hong, S. Lee, J. Burford, G. Gyarmati, J. Peti-Peterdi, S. Srikanth, Y. Gwack, C. J. Koh, E. Boriushkin, A. Hamik, A. K. Wong and Y. K. Hong (2017). “ORAI1 Activates Proliferation of Lymphatic Endothelial Cells in Response to Laminar Flow Through Kruppel-Like Factors 2 and 4.” Circ Res 120(9): 1426–1439.

Chubinskiy-Nadezhdin, V., S. Semenova, V. Vasileva, A. Shatrova, N. Pugovkina and Y. Negulyaev (2022). “Store-Operated Ca(2+) Entry Contributes to Piezo1-Induced Ca(2+) Increase in Human Endometrial Stem Cells.” Int J Mol Sci 23(7).

Cooper, S. T. and P. L. McNeil (2015). “Membrane Repair: Mechanisms and Pathophysiology.” Physiol Rev 95(4): 1205–1240.

Corrotte, M., P. E. Almeida, C. Tam, T. Castro-Gomes, M. C. Fernandes, B. A. Millis, M. Cortez, H. Miller, W. Song, T. K. Maugel and N. W. Andrews (2013). “Caveolae internalization repairs wounded cells and muscle fibers.” Elife 2: e00926.

Corrotte, M., M. C. Fernandes, C. Tam and N. W. Andrews (2012). “Toxin pores endocytosed during plasma membrane repair traffic into the lumen of MVBs for degradation.” Traffic 13(3): 483–494.

Demonbreun, A. R. and E. M. McNally (2016). “Plasma Membrane Repair in Health and Disease.” Curr Top Membr 77: 67–96.

Demonbreun, A. R., M. Quattrocelli, D. Y. Barefield, M. V. Allen, K. E. Swanson and E. M. McNally (2016). “An actin-dependent annexin complex mediates plasma membrane repair in muscle.” J Cell Biol 213(6): 705–718.

Ding, J., J. R. Zhang, Y. Wang, C. L. Li, D. Lu, S. M. Guan and J. Chen (2012). “Effects of a non-selective TRPC channel blocker, SKF-96365, on melittin-induced spontaneous persistent nociception and inflammatory pain hypersensitivity.” Neurosci Bull 28(2): 173–181.

Dupont, S., L. Morsut, M. Aragona, E. Enzo, S. Giulitti, M. Cordenonsi, F. Zanconato, J. Le Digabel, M. Forcato, S. Bicciato, N. Elvassore and S. Piccolo (2011). “Role of YAP/TAZ in mechanotransduction.” Nature 474(7350): 179–183.

Dynes, J. L., A. Amcheslavsky and M. D. Cahalan (2016). “Genetically targeted single-channel optical recording reveals multiple Orai1 gating states and oscillations in calcium influx.” Proc Natl Acad Sci U S A 113(2): 440–445.

Elaib, Z., F. Saller and R. Bobe (2016). “The Calcium Entry-Calcium Refilling Coupling.” Adv Exp Med Biol 898: 333–352.

Fu, R., W. Wang, Y. Huo, L. Li, R. Chen, Z. Lin, Y. Tao, X. Peng, W. Huang and C. Guo (2024). “The mechanosensitive ion channel Piezo1 contributes to podocyte cytoskeleton remodeling and development of proteinuria in lupus nephritis.” Kidney Int 106(4): 625–639.

Garcia, J., J. Bagwell, B. Njaine, J. Norman, D. S. Levic, S. Wopat, S. E. Miller, X. Liu, J. W. Locasale, D. Y. R. Stainier and M. Bagnat (2017). “Sheath Cell Invasion and Trans-differentiation Repair Mechanical Damage Caused by Loss of Caveolae in the Zebrafish Notochord.” Curr Biol 27(13): 1982–1989 e1983.

Gounou, C., F. Bouvet, B. Liet, V. Prouzet-Mauleon, L. d’Agata, E. Harte, F. Argoul, G. Siegfried, R. Iggo, A. M. Khatib and A. Bouter (2023). “Annexin-A5 and annexin-A6 silencing prevents metastasis of breast cancer cells in zebrafish.” Biol Cell 115(6): e202200110.

Hagenbeek, T. J., J. R. Zbieg, M. Hafner, R. Mroue, J. A. Lacap, N. M. Sodir, C. L. Noland, S. Afghani, A. Kishore, K. P. Bhat, X. Yao, S. Schmidt, S. Clausen, M. Steffek, W. Lee, P. Beroza, S. Martin, E. Lin, R. Fong, P. Di Lello, M. H. Kubala, M. N. Yang, J. T. Lau, E. Chan, A. Arrazate, L. An, E. Levy, M. N. Lorenzo, H. J. Lee, T. H. Pham, Z. Modrusan, R. Zang, Y. C. Chen, M. Kabza, M. Ahmed, J. Li, M. T. Chang, D. Maddalo, M. Evangelista, X. Ye, J. J. Crawford and A. Dey (2023). “An allosteric pan-TEAD inhibitor blocks oncogenic YAP/TAZ signaling and overcomes KRAS G12C inhibitor resistance.” Nat Cancer 4(6): 812–828.

Hannezo, E. and C. P. Heisenberg (2019). “Mechanochemical Feedback Loops in Development and Disease.” Cell 178(1): 12–25.

Higashida, C., T. Kiuchi, Y. Akiba, H. Mizuno, M. Maruoka, S. Narumiya, K. Mizuno and N. Watanabe (2013). “F- and G-actin homeostasis regulates mechanosensitive actin nucleation by formins.” Nat Cell Biol 15(4): 395–405.

Hoernke, M., J. Mohan, E. Larsson, J. Blomberg, D. Kahra, S. Westenhoff, C. Schwieger and R. Lundmark (2017). “EHD2 restrains dynamics of caveolae by an ATP-dependent, membrane-bound, open conformation.” Proc Natl Acad Sci U S A 114(22): E4360–e4369.

Horn, A. and J. K. Jaiswal (2018). “Cellular mechanisms and signals that coordinate plasma membrane repair.” Cell Mol Life Sci 75(20): 3751–3770.

Jaiswal, J. K., S. P. Lauritzen, L. Scheffer, M. Sakaguchi, J. Bunkenborg, S. M. Simon, T. Kallunki, M. Jaattela and J. Nylandsted (2014). “S100A11 is required for efficient plasma membrane repair and survival of invasive cancer cells.” Nat Commun 5: 3795.

Jardin, I. and J. A. Rosado (2016). “STIM and calcium channel complexes in cancer.” Biochim Biophys Acta 1863(6 Pt B): 1418–1426.

Lachowski, D., C. Matellan, S. Gopal, E. Cortes, B. K. Robinson, A. Saiani, A. F. Miller, M. M. Stevens and A. E. Del Rio Hernandez (2022). “Substrate Stiffness-Driven Membrane Tension Modulates Vesicular Trafficking via Caveolin-1.” ACS Nano 16(3): 4322–4337.

Lakk, M. and D. Krizaj (2021). “TRPV4-Rho signaling drives cytoskeletal and focal adhesion remodeling in trabecular meshwork cells.” Am J Physiol Cell Physiol 320(6): C1013–C1030.

Lee, J. H., Y. G. Yun and H. W. Kim (2025). “Matrix-induced nuclear remodeling and mechano-therapeutics.” Cell Rep 44(9): 116176.

Leung, C., C. Yu, M. I. Lin, C. Tognon and P. Bernatchez (2013). “Expression of myoferlin in human and murine carcinoma tumors: role in membrane repair, cell proliferation, and tumorigenesis.” Am J Pathol 182(5): 1900–1909.

Lewis, R. S. (2020). “Store-Operated Calcium Channels: From Function to Structure and Back Again.” Cold Spring Harb Perspect Biol 12(5).

Lolo, F. N., N. Walani, E. Seemann, D. Zalvidea, D. M. Pavon, G. Cojoc, M. Zamai, C. Viaris de Lesegno, F. Martinez de Benito, M. Sanchez-Alvarez, J. J. Uriarte, A. Echarri, D. Jimenez-Carretero, J. C. Escolano, S. A. Sanchez, V. R. Caiolfa, D. Navajas, X. Trepat, J. Guck, C. Lamaze, P. Roca-Cusachs, M. M. Kessels, B. Qualmann, M. Arroyo and M. A. Del Pozo (2023). “Caveolin-1 dolines form a distinct and rapid caveolae-independent mechanoadaptation system.” Nat Cell Biol 25(1): 120–133.

Luan, H., T. A. Bielecki, B. C. Mohapatra, N. Islam, I. Mushtaq, A. M. Bhat, S. Mirza, S. Chakraborty, M. Raza, M. D. Storck, M. S. Toss, J. L. Meza, W. B. Thoreson, D. W. Coulter, E. A. Rakha, V. Band and H. Band (2023). “EHD2 overexpression promotes tumorigenesis and metastasis in triple-negative breast cancer by regulating store-operated calcium entry.” Elife 12.

Marg, A., V. Schoewel, T. Timmel, A. Schulze, C. Shah, O. Daumke and S. Spuler (2012). “Sarcolemmal repair is a slow process and includes EHD2.” Traffic 13(9): 1286–1294.

Martino, F., A. R. Perestrelo, V. Vinarsky, S. Pagliari and G. Forte (2018). “Cellular Mechanotransduction: From Tension to Function.” Front Physiol 9: 824.

McMahon, H. T. and E. Boucrot (2015). “Membrane curvature at a glance.” J Cell Sci 128(6): 1065–1070.

Miller, J., C. Bruen, M. Schnaus, J. Zhang, S. Ali, A. Lind, Z. Stoecker, K. Stauderman and S. Hebbar (2020). “Auxora versus standard of care for the treatment of severe or critical COVID-19 pneumonia: results from a randomized controlled trial.” Crit Care 24(1): 502.

Miroshnikova, Y. A., S. Manet, X. Li, S. A. Wickstrom, E. Faurobert and C. Albiges-Rizo (2021). “Calcium signaling mediates a biphasic mechanoadaptive response of endothelial cells to cyclic mechanical stretch.” Mol Biol Cell 32(18): 1724–1736.

Moren, B., C. Shah, M. T. Howes, N. L. Schieber, H. T. McMahon, R. G. Parton, O. Daumke and R. Lundmark (2012). “EHD2 regulates caveolar dynamics via ATP-driven targeting and oligomerization.” Mol Biol Cell 23(7): 1316–1329.

Moreno-Vicente, R., D. M. Pavon, I. Martin-Padura, M. Catala-Montoro, A. Diez-Sanchez, A. Quilez-Alvarez, J. A. Lopez, M. Sanchez-Alvarez, J. Vazquez, R. Strippoli and M. A. Del Pozo (2018). “Caveolin-1 Modulates Mechanotransduction Responses to Substrate Stiffness through Actin-Dependent Control of YAP.” Cell Rep 25(6): 1622–1635 e1626.

Narain, R., J. M. Muncie-Vasic and V. M. Weaver (2025). “Forcing the code: tension modulates signaling to drive morphogenesis and malignancy.” Genes Dev 39(1-2): 163–181.

Nieto-Felipe, J., A. Macias-Diaz, J. Sanchez-Collado, A. Berna-Erro, I. Jardin, G. M. Salido, J. J. Lopez and J. A. Rosado (2023). “Role of Orai-family channels in the activation and regulation of transcriptional activity.” J Cell Physiol 238(4): 714–726.

Ong, H. L., K. P. Subedi, G. Y. Son, X. Liu and I. S. Ambudkar (2019). “Tuning store-operated calcium entry to modulate Ca(2+)-dependent physiological processes.” Biochim Biophys Acta Mol Cell Res 1866(7): 1037–1045.

Pallagi, P., M. Görög, N. Papp, T. Madácsy, Á. Varga, T. Crul, V. Szabó, M. Molnár, K. Dudás, A. Grassalkovich, E. Szederkényi, G. Lázár, V. Venglovecz, P. Hegyi and J. Maléth (2022). “Bile acid- and ethanol-mediated activation of Orai1 damages pancreatic ductal secretion in acute pancreatitis.” J Physiol 600(7): 1631–1650.

Pardo-Pastor, C., F. Rubio-Moscardo, M. Vogel-Gonzalez, S. A. Serra, A. Afthinos, S. Mrkonjic, O. Destaing, J. F. Abenza, J. M. Fernandez-Fernandez, X. Trepat, C. Albiges-Rizo, K. Konstantopoulos and M. A. Valverde (2018). “Piezo2 channel regulates RhoA and actin cytoskeleton to promote cell mechanobiological responses.” Proc Natl Acad Sci U S A 115(8): 1925–1930.

Parsonage, G., K. Cuthbertson, N. Endesh, N. Murciano, A. J. Hyman, C. H. Revill, O. V. Povstyan, E. Chuntharpursat-Bon, M. Debant, M. J. Ludlow, T. S. Futers, L. Lichtenstein, J. A. Kinsella, F. Bartoli, M. G. Rotordam, N. Becker, A. Bruggemann, R. Foster and D. J. Beech (2023). “Improved PIEZO1 agonism through 4-benzoic acid modification of Yoda1.” Br J Pharmacol 180(16): 2039–2063.

Parton, R. G. (2018). “Caveolae: Structure, Function, and Relationship to Disease.” Annu Rev Cell Dev Biol 34: 111–136.

Posey, A. D., Jr., P. Pytel, K. Gardikiotes, A. R. Demonbreun, M. Rainey, M. George, H. Band and E. M. McNally (2011). “Endocytic recycling proteins EHD1 and EHD2 interact with fer-1-like-5 (Fer1L5) and mediate myoblast fusion.” J Biol Chem 286(9): 7379–7388.

Ramsey, I. S., M. Delling and D. E. Clapham (2006). “An introduction to TRP channels.” Annu Rev Physiol 68: 619–647.

Rausch, V., J. R. Bostrom, J. Park, I. R. Bravo, Y. Feng, D. C. Hay, B. A. Link and C. G. Hansen (2019). “The Hippo Pathway Regulates Caveolae Expression and Mediates Flow Response via Caveolae.” Curr Biol 29(2): 242–255 e246.

Sathish, V., A. J. Abcejo, M. A. Thompson, G. C. Sieck, Y. S. Prakash and C. M. Pabelick (2012). “Caveolin-1 regulation of store-operated Ca(2+) influx in human airway smooth muscle.” Eur Respir J 40(2): 470–478.

Shvets, E., A. Ludwig and B. J. Nichols (2014). “News from the caves: update on the structure and function of caveolae.” Curr Opin Cell Biol 29: 99–106.

Sinha, B., D. Koster, R. Ruez, P. Gonnord, M. Bastiani, D. Abankwa, R. V. Stan, G. Butler-Browne, B. Vedie, L. Johannes, N. Morone, R. G. Parton, G. Raposo, P. Sens, C. Lamaze and P. Nassoy (2011). “Cells respond to mechanical stress by rapid disassembly of caveolae.” Cell 144(3): 402–413.

Sotodosos-Alonso, L., M. Pulgarin-Alfaro and M. A. Del Pozo (2023). “Caveolae Mechanotransduction at the Interface between Cytoskeleton and Extracellular Matrix.” Cells 12(6).

Stauderman, K. A. (2018). “CRAC channels as targets for drug discovery and development.” Cell Calcium 74: 147–159.

Stefl, M., M. Takamiya, V. Middel, M. Tekpinar, K. Nienhaus, T. Beil, S. Rastegar, U. Strahle and G. U. Nienhaus (2024). “Caveolae disassemble upon membrane lesioning and foster cell survival.” iScience 27(2): 108849.

Stoeber, M., I. K. Stoeck, C. Hanni, C. K. Bleck, G. Balistreri and A. Helenius (2012). “Oligomers of the ATPase EHD2 confine caveolae to the plasma membrane through association with actin.” EMBO J 31(10): 2350–2364.

Suchyna, T. M., J. H. Johnson, K. Hamer, J. F. Leykam, D. A. Gage, H. F. Clemo, C. M. Baumgarten and F. Sachs (2000). “Identification of a peptide toxin from Grammostola spatulata spider venom that blocks cation-selective stretch-activated channels.” J Gen Physiol 115(5): 583–598.

Szabo, V., N. Csakany-Papp, M. Gorog, T. Madacsy, A. Varga, A. Kiss, B. Tel, B. Jojart, T. Crul, K. Dudas, M. Bagyanszki, N. Bodi, F. Ayaydin, S. Jee, L. Tiszlavicz, K. A. Stauderman, S. Hebbar, P. Pallagi and J. Maleth (2023). “Orai1 calcium channel inhibition prevents progression of chronic pancreatitis.” JCI Insight 8(13).

Szabó, V., N. Csákány-Papp, M. Görög, T. Madacsy, Á. Varga, A. Kiss, B. Tel, B. Jójárt, T. Crul, K. Dudás, M. Bagyánszki, N. Bódi, F. Ayaydin, S. Jee, L. Tiszlavicz, K. A. Stauderman, S. Hebbar, P. Pallagi and J. Maléth (2023). “Orai1 calcium channel inhibition prevents progression of chronic pancreatitis.” JCI Insight.

Torrino, S., W. W. Shen, C. M. Blouin, S. K. Mani, C. Viaris de Lesegno, P. Bost, A. Grassart, D. Koster, C. A. Valades-Cruz, V. Chambon, L. Johannes, P. Pierobon, V. Soumelis, C. Coirault, S. Vassilopoulos and C. Lamaze (2018). “EHD2 is a mechanotransducer connecting caveolae dynamics with gene transcription.” J Cell Biol 217(12): 4092–4105.

van Vliet, A. R., F. Giordano, S. Gerlo, I. Segura, S. Van Eygen, G. Molenberghs, S. Rocha, A. Houcine, R. Derua, T. Verfaillie, J. Vangindertael, H. De Keersmaecker, E. Waelkens, J. Tavernier, J. Hofkens, W. Annaert, P. Carmeliet, A. Samali, H. Mizuno and P. Agostinis (2017). “The ER Stress Sensor PERK Coordinates ER-Plasma Membrane Contact Site Formation through Interaction with Filamin-A and F-Actin Remodeling.” Mol Cell 65(5): 885–899.e886.

Varadarajan, S., S. A. Chumki, R. E. Stephenson, E. R. Misterovich, J. L. Wu, C. E. Dudley, I. S. Erofeev, A. B. Goryachev and A. L. Miller (2022). “Mechanosensitive calcium flashes promote sustained RhoA activation during tight junction remodeling.” J Cell Biol 221(4).

Waldron, R. T., Y. Chen, H. Pham, A. Go, H. Y. Su, C. Hu, L. Wen, S. Z. Husain, C. A. Sugar, J. Roos, S. Ramos, A. Lugea, M. Dunn, K. Stauderman and S. J. Pandol (2019). “The Orai Ca(2+) channel inhibitor CM4620 targets both parenchymal and immune cells to reduce inflammation in experimental acute pancreatitis.” J Physiol 597(12): 3085–3105.

Wei, Y. and W. Li (2021). “Calcium, an Emerging Intracellular Messenger for the Hippo Pathway Regulation.” Front Cell Dev Biol 9: 694828.

Wu, J., L. Liu, T. Matsuda, Y. Zhao, A. Rebane, M. Drobizhev, Y. F. Chang, S. Araki, Y. Arai, K. March, T. E. Hughes, K. Sagou, T. Miyata, T. Nagai, W. H. Li and R. E. Campbell (2013). “Improved orange and red Ca(2)+/- indicators and photophysical considerations for optogenetic applications.” ACS Chem Neurosci 4(6): 963–972.

Yang, S., X. Miao, S. Arnold, B. Li, A. T. Ly, H. Wang, M. Wang, X. Guo, M. M. Pathak, W. Zhao, C. D. Cox and Z. Shi (2022). “Membrane curvature governs the distribution of Piezo1 in live cells.” Nat Commun 13(1): 7467.

Yang, Y., L. A. Valencia, C. H. Lu, M. L. Nakamoto, C. T. Tsai, C. Liu, H. Yang, W. Zhang, Z. Jahed, W. R. Lee, F. Santoro, J. Liou, J. C. Wu and B. Cui (2024). “Plasma membrane curvature regulates the formation of contacts with the endoplasmic reticulum.” Nat Cell Biol 26(11): 1878–1891.

Yeow, I., G. Howard, J. Chadwick, C. Mendoza-Topaz, C. G. Hansen, B. J. Nichols and E. Shvets (2017). “EHD Proteins Cooperate to Generate Caveolar Clusters and to Maintain Caveolae during Repeated Mechanical Stress.” Curr Biol 27(19): 2951–2962 e2955.

Yeow, I., G. Howard, J. Chadwick, C. Mendoza-Topaz, C. G. Hansen, B. J. Nichols and E. Shvets (2017). “EHD Proteins Cooperate to Generate Caveolar Clusters and to Maintain Caveolae during Repeated Mechanical Stress.” Curr Biol 27(19): 2951–2962.e2955.

