## Supplementary figure legends for "Orai1 is required for Ca^2+^-dependent plasma membrane repair and mechanoadaptation"

**Figure S1. Related to Figure 1.** Representative FITC-Dextran (green; top), propidium iodide (red; middle) and merged (bottom) fluorescence images of mechanically-induced plasma membrane repair in MDA-MB-231 (**S1A**) and Hs578T (**S1B**) cell lines 20 min post plasma membrane damage. Scale bars, 200 µm.

**Figure S2. Related to Figure 2.** **S2A.** Migration assay results of wildtype (WT) vs. ORAI1-KO MDA-MB-231 and Hs578T cell lines. **S2B.** Representative FITC-Dextran (green; left), propidium iodide (PI, red; middle) and merged (right) fluorescence images of the repair of mechanically induced plasma membrane damage in MDA-MB-231 and Hs578T cell lines at 20-minute time point. **S2C**. Migration assay results of wildtype (WT) and ORAI1-KO MDA-MB-231 cell line treated with vehicle (DMSO) or CM4620 (10 μM; 18 hours). **S2D.** Representative FITC-Dextran (green; left), propidium iodide (PI, red; middle) and merged (right) fluorescence images of the repair of mechanically-induced plasma membrane damage in MDA-MB-231 cell line at 20-minute time point after 72-hour pretreatment with DMSO (control) or CM4620 (10 μM). Scale bars, 200 µm.

**Figure S3. Related to Figure 4. S3A.** Thapsigargin (TG)-induced SOCE measurements (initial peak, Ca^2+^ store release in the absence of extracellular Ca^2+^; second peak, SOCE in the presence of extracellular Ca^2+^) performed on the indicated cell lines cultured with or without CM5480 (10 μM, 4 hours pretreatment) (Left panel). Migration assay results of wildtype (WT) and ORAI1-KO MDA-MB-231 cell line treated with vehicle (DMSO) or CM5480 (10 μM; 18 hours) (Right panel). **S3B.** Inhibition of SOCE by CM5480. Thapsigargin (TG)-induced SOCE measurements (initial peak, Ca^2+^ store release in the absence of extracellular Ca^2+^; second peak, SOCE in the presence of extracellular Ca^2+^) performed on MDA-MB-231 transfected with Orai1 fused to a Ca^2+^ indicator (G-GECO1-Orai1) and cultured with or without CM5480 (10 μM, 4 hours pretreatment). **S3C.** Anti-STIM1 or Anti-STIM2 Western blotting of MDA-MB-231 cells transfected with various amounts (picomoles, pmol) of STIM1 or STIM2 siRNA. β-actin, loading control. **S3D.** Anti-STIM1 or Anti-STIM2 Western blotting of MDA-MB-231 cells co-transfected with STIM1 and STIM2 siRNA (100 pmol. each); β-actin, loading control.

**Figure S4.** **Related to Figure 5.** **A.** Representative confocal images of YAP (green; left), DAPI (blue; middle) and merged (right) staining of MDA-MB-231 cell lines of the indicated genotypes cultured on 0.2 kPa or 64 kPa hydrogels. **B.** Representative Confocal images of YAP (green; left), DAPI (blue; middle) and merged (right) staining in MDA-MB-231 cell lines following treatment with control (DMSO), CM5480 (10 μM) or CM4620 (10 μM) for 48 hours.

**Figure S5.** **Related to Figure 6.** Representative FITC-Dextran (green; left), propidium iodide (PI, red; middle) and merged (right) fluorescence images of the repair of mechanically-induced plasma membrane damage in MDA-MB-231 and Hs578T cell lines at 20-minute time points after 24-hour pretreatment without (control) or with GNE-7883 (10 μM).

**Video S1. Time-lapse imaging of an MDA-MB-231 cell expressing EHD2-mCherry (red) to visualize the response to localized mechanical indentation with a microcantilever; related to Figure 3.**

**Video S2.** **Time-lapse imaging of MDA-MB-231 WT, EHD2-KO, ORAI1-KO, CAV1-KO, and mouse-EHD2-reconstituted EHD2-KO cells expressing R-GECO1.2 (red) to visualize the influx of Ca^2+^ upon localized mechanical indentation with a microcantilever; related to Figure 4B.**

**Video S3.** **Time-lapse imaging of MDA-MB-231 expressing G-GECO1-Orai1 (green) to visualize the influx of Ca^2+^ upon localized mechanical indentation with a microcantilever; related to Figure 4D.**
